# Likelihood-Free Parameter Inference for Spatiotemporal Stochastic Biological Models using Neural Posterior Estimation

**DOI:** 10.1101/2025.10.26.684706

**Authors:** Tom Kimpson, Jennifer Flegg, Matthew J. Simpson

## Abstract

Cell migration is a key biological process underlying wound healing, tissue development, and cancer metastasis, yet calibrating mathematical models of migration to experimental data remains a major challenge. Scratch and barrier assays are widely used to study collective cell spreading, and agent-based random walk models provide a natural stochastic description of these experiments. However, parameter inference for such models is hampered by intractable likelihoods, forcing researchers to rely on Approximate Bayesian Computation, which introduces biases and tuning difficulties, or surrogate models that require potentially erroneous noise model specifications. Here, we overcome these limitations using neural posterior estimation, a simulation-based inference framework that learns the full posterior distribution directly from stochastic simulations without surrogate approximations or explicit noise model specifications. We deploy this framework on four progressively complex random walk models of barrier assay experiments describing in vitro cell migration: an isotropic baseline, a model with directional bias (chemotaxis), a model with cell proliferation, and a combined model incorporating both bias and proliferation. For each model, we demonstrate inference in two settings: using one-dimensional summary statistics (column counts), and using a convolutional neural network that enables inference directly from raw two-dimensional spatial data. Neural posterior estimation performs well across all four models, recovering biologically interpretable parameters (e.g. cell motility, directional bias, proliferation rates) from cases where classical surrogate-based methods are adequate through to the combined model where the interplay of multiple mechanisms renders surrogate approximations unreliable. We validate all posteriors using simulation-based calibration diagnostics and provide an open-source implementation of our pipeline to facilitate its adoption and extension to more complex, spatially-structured biological models.

## 1 Introduction

Scratch and barrier assay experiments are widely used to study how cell populations spread, proliferate, and interact in vitro [1, 2, 3], with applications spanning wound healing, embryonic development, immune responses, and cancer metastasis [4, 5]. In these experiments, cells are confined to a region on a substrate; after barrier removal, the population spreads outward, and the resulting spatiotemporal dynamics encode information about the underlying motility and proliferation mechanisms. Extracting this information quantitatively requires calibrating mathematical models to experimental data, yet doing so remains a major challenge for stochastic, agent-based descriptions of cell migration.

Agent-based random walk models on regular lattices provide a natural stochastic description of these experiments, capturing the inherent biological variability and heterogeneity that deterministic continuum models cannot adequately represent [6, 7, 8]. However, calibrating these discrete stochastic models to data poses serious inferential challenges. The likelihood function is typically intractable, precluding standard statistical methods such as Maximum Likelihood Estimation (MLE) and Markov chain Monte Carlo (MCMC). Classical likelihood-free alternatives like Approximate Bayesian Computation (ABC) are computationally expensive and sensitive to tuning choices [9, 10]. A common workaround is to replace the stochastic simulator with a deterministic partial differential equation (PDE) surrogate, but this introduces systematic bias and requires specifying an explicit noise model whose misspecification can lead to poorly calibrated uncertainties [11, 12, 13]. These practical difficulties have motivated a growing interest in simulation-based inference methods that can learn directly from stochastic simulators [14, 15, 7, 16, 17, 18].

In this paper, we address these challenges using Neural Posterior Estimation (NPE), a simulation-based inference method that learns the posterior distribution *p*(*θ* |*x*) directly from parameter-data pairs generated by the stochastic simulator [19, 20, 21]. NPE is likelihood-free, learns directly from the high-fidelity simulator without surrogate approximations, and implicitly captures the stochastic relationship between parameters and data without an explicit noise model. NPE also provides amortized inference, where the upfront cost of training is offset by near-instant posterior estimation for any new observation. A further key aspect of our approach is the integration of a Convolutional Neural Network (CNN) as a learned feature extractor within the NPE pipeline. In standard practice, two-dimensional spatial data from barrier assays are reduced to one-dimensional summary statistics such as column-averaged cell counts, potentially discarding informative spatial structure. By embedding a CNN directly into the inference framework, we enable posterior estimation from full two-dimensional lattice configurations without hand-crafted summary statistics, allowing the network to automatically learn which spatial features are most informative for parameter estimation.

We demonstrate this framework on a hierarchy of agent-based random walk models of a barrier assay experiment, following the framework of [7] (S&P2025 hereafter) as our reference. Beginning with an isotropic baseline, we progressively add directional bias (chemotaxis), cell proliferation, and their combination. This hierarchy illustrates a progression from models where well-validated surrogate likelihoods exist and NPE is a convenient alternative to classical methods, to a combined model where the interplay of multiple mechanisms renders mean-field surrogate approximations increasingly unreliable and simulation-based inference provides a principled route to full Bayesian inference. The remainder of this paper is organized as follows. Section 2 describes the random walk models, reviews classical inference approaches and their limitations, and details the NPE methodology and our CNN-based data processing pipeline. Section 3 presents inference results across all four models using both one-dimensional summary statistics and raw two-dimensional spatial data, validates posteriors using simulation-based calibration diagnostics, and compares computational costs. Section 4 discusses the implications of our findings for simulation-based inference in biology.

## 2 Methods

### 2.1 Random Walk Model

We study a hierarchy of stochastic agent-based models of increasing complexity. Here we describe the baseline model; extensions incorporating directional bias, proliferation, and their combination are introduced progressively in Section 3. Agents representing biological cells are initially confined to a central region on a two-dimensional square lattice and subsequently spread outwards via an unbiased random walk. The experimental setup, illustrated in Figure 1, mimics a barrier assay where cells are placed in a confined region and allowed to spread after barrier removal. The two-dimensional lattice has height *H* = 50 (with 0≤ *y* ≤ 50) and width *W* = 200 (with − 100 ≤ *x*≤ 100), with lattice spacing Δ = 1 and discrete time step *τ* = 1. Zero-flux boundary conditions are applied at all edges. Initial conditions are set by occupying each site in the central region| *x* |≤ 25 with a single agent with probability *U* = 0.3, as shown in Figure 1a. At each time step, the system evolves through a random sequential update method where agents move to one of four nearest-neighbour sites with probability *P* = 0.7. After 100 time steps, the population spreads symmetrically as shown in Figure 1b.

**Figure 1.**
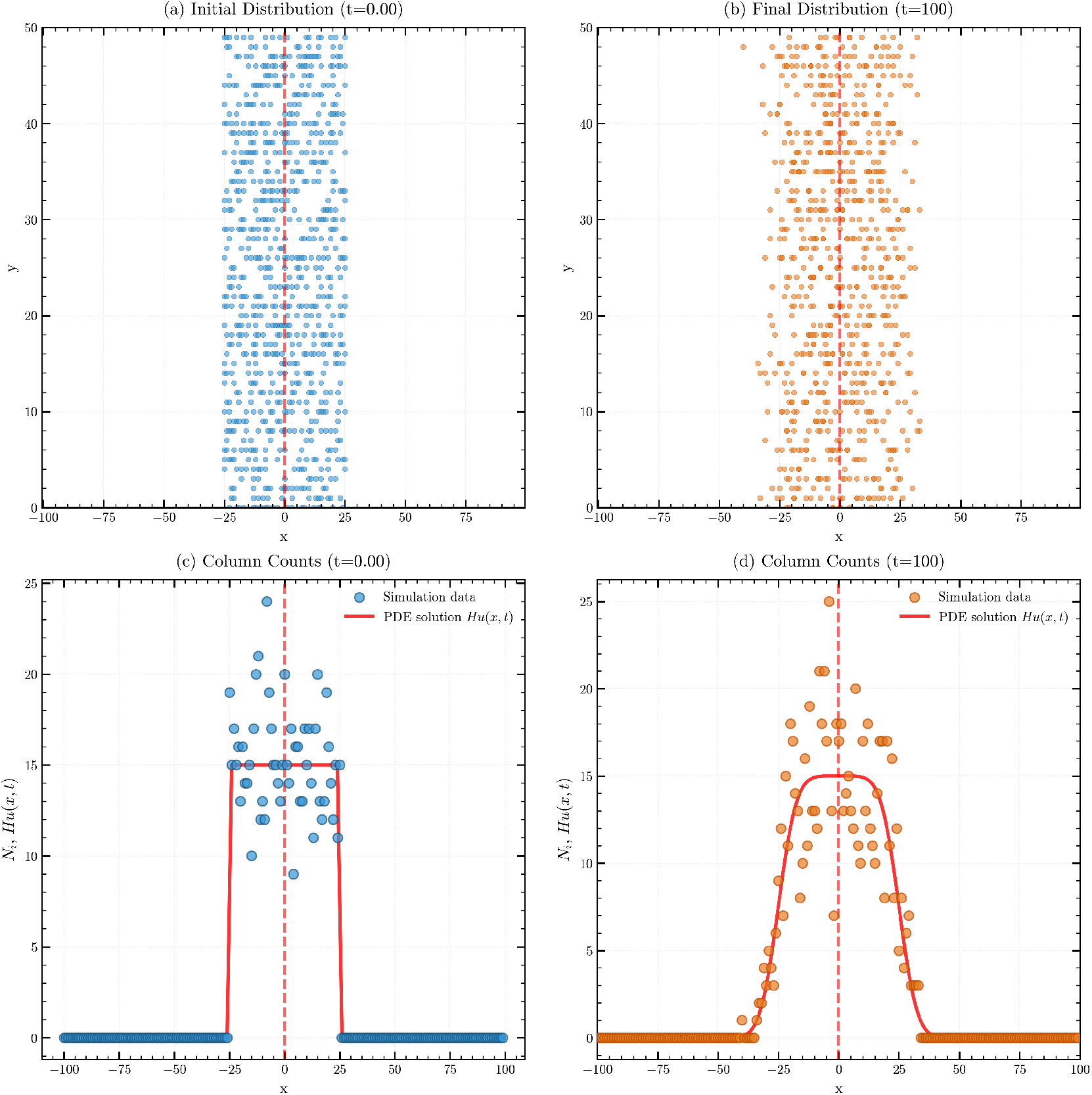
Schematic of the lattice-based random walk model. (a) Initial placement of agents within region| *x*|≤ 25 on a 2D lattice with *H* = 50, Δ = *τ* = 1, *U* = 0.3. (b) Agent distribution after 100 time steps showing symmetric outward spreading. (c)–(d) Reduction of 2D data to 1D column counts *N*_*i*_(*t*) at *t* = 0 and *t* = 100 respectively. Blue dots show initial stochastic count data; orange dots show final stochastic count data; red curves show the continuum PDE solution *Hu*(*x, t*) where *H* is the lattice height and *u*(*x, t*) solves the diffusion equation with *D* = *P/*4.

For lattice-based random walk models of this type, the mean behaviour of agents in the continuum limit can be described by a PDE of the form [6, 7, 8]

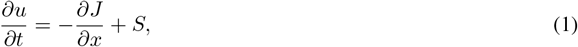

where *u*(*x, t*) represents mean agent density, *J* is the macroscopic flux, and *S* denotes source terms from proliferation. For the simple case considered here (unbiased motion, no proliferation, no crowding), this reduces to the linear diffusion equation

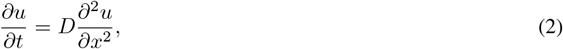

where *D* = lim_Δ,*τ→*0_ *P* Δ^2^*/*(4*τ* ) is the effective diffusion coefficient, subject to zero-flux boundary conditions*∂u/∂x* = 0 at *x* = *±*100 and initial condition *u*(*x*, 0) = *U* for |*x*| ≤ 25, *u*(*x*, 0) = 0 otherwise. Working with lattice dimensionless units Δ = *τ* = 1, this gives *D ≈ P/*4 = 0.175 for our simulation with *P* = 0.7. The two-dimensional data can also be reduced to one-dimensional column counts *N*_*i*_(*t*) = Σ_*j*_ **1**[site (*i, j*) occupied], as illustrated in Figure 1c–d, where the continuum PDE solution *Hu*(*x, t*) provides a good approximation to the mean column counts.

While S&P2025 investigates a suite of model variations and alternative PDE approximations, we use this unbiased, non-interacting case as our baseline (the “original model”) and progressively extend the analysis to three additional models. Model A introduces directional bias via a parameter *ρ* controlling the probability of stepping in the positive *x*-direction. Model B incorporates cell proliferation through a Fisher–Kolmogorov mechanism with rate *R*. Model C combines both bias and proliferation, representing the most biologically realistic scenario. We refer the reader to S&P2025 for full model specifications and focus here on demonstrating how NPE handles the increasing complexity across this model hierarchy.

### 2.2 Classical Inference Approaches

We briefly review the established approaches to inference for stochastic agent-based models, drawing primarily on the treatment in S&P2025, which provides systematic comparisons across multiple inference paradigms for biological random-walk models. Table 1 provides a comparative overview.

**Table 1:** Comparison of inference approaches for stochastic biological models. ^*^The low per-observation cost for NPE is achieved after an initial, computationally intensive training phase. ^†^Amortized methods incur a one-time training cost after which per-observation cost is negligible.

| Method | Per-observation Cost | Requires Surrogate? | Requires Summary Stats? | Amortized?<sup>†</sup> |
| --- | --- | --- | --- | --- |
| ABC | Very High | No | Yes | No |
| Surrogate + MCMC | High | Yes | Yes | No |
| Surrogate + MLE | Medium | Yes | Yes | No |
| **NPE (proposed)** | **Low<sup>\*</sup>** | **No** | **Optional** | **Yes** |

#### 2.2.1 Approximate Bayesian Computation

ABC represents the most direct approach to likelihood-free inference for stochastic models [9, 10, 22, 23]. The foundational ABC rejection sampling algorithm approximates the posterior distribution by undertaking the following steps: (i) a parameter vector *θ*^(*j*)^ is drawn from the prior distribution *p*(*θ*); (ii) a dataset *Y* ^(*j*)^ is simulated from the model using *θ*^(*j*)^; (iii) a discrepancy between the summary statistics of the observed data, *s*(*Y*_obs_), and simulated data, *s*(*Y* ^(*j*)^), is calculated using a distance metric *ρ*(*·,·*); and (iv) the parameter *θ*^(*j*)^ is accepted if this discrepancy is less than a tolerance threshold, *ρ*(*s*(*Y*_obs_), *s*(*Y* ^(*j*)^)) ≤ *ϵ*. The collection of accepted parameters forms a sample from an approximation to the true posterior, *p*(*θ* | *Y*_obs_) [24].

Despite its conceptual simplicity, ABC faces significant methodological and computational challenges. A primary issue is the choice of the tolerance *ϵ*, which defines a fundamental bias-accuracy trade-off [9, 25]. Only in the idealised limit where *ϵ→* 0 and the summary statistics are sufficient does the ABC posterior converge to the true posterior [26, 25]. In practice, a smaller *ϵ* reduces bias but dramatically lowers the acceptance rate, increasing computational cost. Conversely, a larger *ϵ* increases the acceptance rate at the cost of yielding a more biased and diffuse posterior approximation.

This trade-off is exacerbated by the “curse of dimensionality” [27, 25]. For models with high-dimensional data or parameter spaces, the acceptance rate can become vanishingly small, demanding a large number of simulations. For instance, S&P2025 generate 10^5^ simulations from their computationally expensive random walk model to retain just the top 1%, highlighting this significant computational inefficiency. Furthermore, the method’s performance depends critically on the choice of summary statistics. If the statistics are not formally sufficient, crucial information from the data is discarded, which introduces a form of bias that cannot be removed by simply lowering the tolerance *ϵ* [28, 26]. Selecting informative yet low-dimensional summary statistics in a principled manner remains a difficult and often ad-hoc challenge [29, 25]. More efficient variants, such as Sequential Monte Carlo ABC [30, 31], iteratively refine a sequence of proposal distributions toward the posterior by progressively lowering the tolerance, improving the acceptance rate relative to simple rejection sampling. Nevertheless, the fundamental reliance on summary statistics and repeated simulation remains.

#### 2.2.2 Surrogate Model Approaches

To mitigate the computational burden of methods like ABC, an alternative strategy replaces the stochastic model with a computationally cheaper surrogate model. A common choice is a deterministic PDE that captures the system’s mean-field behaviour, yielding a tractable likelihood function [32, 33] that enables standard, efficient likelihood-based inference algorithms like MCMC. However, this computational gain comes at the cost of introducing potential model misspecification bias, as inference is performed on an approximation of the true stochastic process.

The primary limitation of this approach is that the resulting inference is conditional on the fidelity of the surrogate. Any discrepancy between the surrogate and the true stochastic process introduces a systematic bias that cannot be reduced by collecting more data [11, 12]. While the continuum limit accurately describes the mean behaviour of non-interacting random walks, the mean-field PDE neglects crucial stochastic effects in more complex scenarios. For example, in interacting models with crowding effects discussed in S&P2025 (Section 4), the mean-field approximation captures average behaviour but cannot represent the bounded nature of agent occupancy or the discrete stochastic fluctuations [34, 33] around this mean. This limitation becomes particularly important when constructing prediction intervals, where the choice of noise model (e.g. Gaussian vs. binomial, see Section 2.2.4) significantly affects the physical realism of predictions [35, 36]. These limitations of surrogate-based approaches motivate the need for inference methods that can work directly with the full stochastic simulator.

#### 2.2.3 Inference with Surrogate Likelihoods

A tractable likelihood function from a surrogate model unlocks standard, efficient inference techniques that would be impossible with the original stochastic simulator. These techniques generally fall into two categories: direct optimization or posterior sampling.

The first category, direct optimization, seeks to find the single parameter set that maximizes the likelihood function, yielding the MLE. While computationally fast, the MLE provides only a point estimate and does not inherently quantify parameter uncertainty, although indirect (asymptotic) uncertainty quantification can be obtained through likelihood ratio tests and Wilks’ theorem under standard regularity conditions. The second category, posterior sampling, uses methods like MCMC to approximate the full posterior distribution, offering a complete characterisation of uncertainty. However, MCMC typically requires significant computational effort, careful hyper-parameter tuning, and rigorous convergence assessment. A third approach, the Laplace approximation, offers a compromise between the speed of optimization and the detailed uncertainty characterization of MCMC. This method approximates the posterior distribution as a multivariate normal distribution centred at the posterior mode (which equals the MLE, 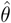 under uniform priors). The covariance is given by the inverse of the observed Fisher Information matrix

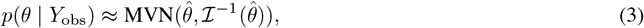

where 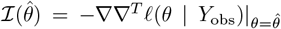 is the negative Hessian of the log-likelihood evaluated at the MLE, often computed using automatic differentiation. While prediction intervals generated via the Laplace approximation can closely match those from more computationally intensive methods (see S&P2025), the method relies on the assumption that the posterior is locally Gaussian. This assumption may be violated for complex models, models with limited data, or simply for models with bounded parameters (e.g., *D≥* 0; see Section 3.1.1) potentially leading to a poor representation of the true uncertainty.

However, all these likelihood-based approaches require an explicit specification of how the deterministic surrogate output relates to the noisy observational data—a critical modelling choice we examine next.

#### 2.2.4 Noise Model Specification

The use of deterministic surrogates necessitates an explicit choice of how to model the relationship between the PDE output and the inherently noisy observations from the stochastic process. This choice profoundly impacts inference results, as the “noise” represents genuine stochastic variability in the agent-based system rather than measurement error.

S&P2025 systematically compare three noise models for relating observed counts to the PDE solution: Gaussian, Poisson, and binomial. The Gaussian noise model assumes observations are normally distributed around the model prediction with constant variance. While mathematically convenient, this leads to unphysical prediction intervals that include negative counts in low-density regions where the population density approaches zero, and fails to respect carrying capacity constraints in high-density regions. The Poisson noise model provides a more natural choice for count data, eliminating the need for an additional variance parameter and ensuring non-negative predictions. The binomial noise model is appropriate for exclusion processes where occupancy is bounded, naturally capturing saturation effects.

S&P2025’s extensive comparisons reveal that noise model misspecification can significantly bias parameter estimates and lead to poorly calibrated prediction intervals. Quantitatively, the Gaussian model consistently produces unphysical negative predictions in low-density regions, while the Poisson and binomial models better respect the discrete, non-negative nature of count data. Similar conclusions regarding the superiority of binomial measurement models over standard additive Gaussian assumptions for biological count data have been demonstrated elsewhere [13].

### 2.3 Neural Posterior Estimation

#### 2.3.1 Overview

NPE [19, 20, 21] takes a distinct approach to inference for simulator-based models within the broader simulation-based inference (SBI) framework. Rather than relying on surrogate approximations or requiring explicit noise models, NPE learns the posterior distribution *p*(*θ* |*Y* ) directly from simulations of the stochastic model. This is achieved through conditional density estimation using neural networks.

While the likelihood *p*(*Y* |*θ*) may be intractable for complex stochastic simulators, we can still readily generate samples from it by running the model simulator. NPE leverages these (parameter, sample) pairs to train a neural network that approximates the inverse mapping—from an observation back to the distribution over parameters that could have produced it. This process is amortized: once the network is trained, it can perform posterior inference for any new observation in seconds rather than hours, without requiring additional simulations. For the class of stochastic biological models considered here, this approach directly addresses each limitation of classical methods: it circumvents intractable likelihoods, eliminates surrogate model bias, implicitly captures complex stochastic relationships without explicit noise models, and can process high-dimensional data through automatic feature extraction.

Figure 2 provides a schematic overview of the NPE workflow, illustrating the distinction between the computationally intensive training phase and the rapid inference phase. During training, parameters are sampled from the prior *p*(*θ*) and used to generate simulated data via the stochastic simulator *p*(*Y* |*θ*). These parameter-data pairs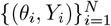 are then used to train a neural density estimator *q*_*ϕ*_(*θ*| *Y* ) that learns to approximate the posterior distribution. Once trained, the inference phase is nearly instantaneous: given new observed data *Y*_obs_, the network directly evaluates the approximate posterior *p*(*θ*|*Y*_obs_) without requiring additional simulations.

**Figure 2.**
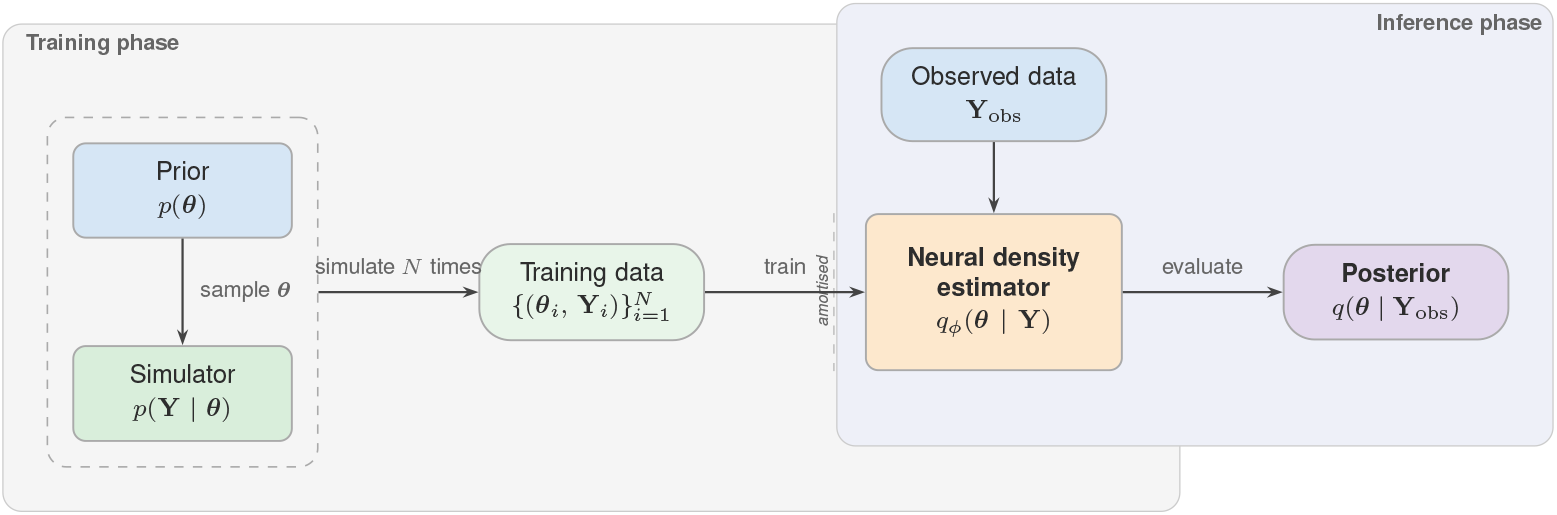
Schematic overview of the NPE workflow. The training phase (left) involves sampling parameters from the prior, generating simulated data, and training a neural density estimator on the resulting parameter-data pairs. The inference phase (right) uses the trained network to instantly evaluate the posterior for new observations without additional simulations.

#### 2.3.2 Conditional Normalizing Flows

At the core of NPE are normalizing flows [37, 38], a class of generative models that construct complex probability distributions by transforming samples from a simple base distribution (typically a standard Gaussian) through a series of invertible, differentiable mappings [39]. Because each mapping is invertible and has a tractable Jacobian, the resulting density can be evaluated exactly via the change-of-variables formula, making normalizing flows well suited to the conditional density estimation task required by NPE.

Formally, let *u ∈* ℝ^*d*^ be a random variable with base density *p*_*u*_(*u*) = 𝒩 (0, *I*), where *I ∈* ℝ^*d×d*^ denotes the identity matrix. The flow defines an invertible transformation *f*_*ϕ*_ : ℝ^*d*^ *→* ℝ^*d*^ parametrized by neural network weights *ϕ*. For conditional density estimation, this transformation is conditioned on the observation: *θ* = *f*_*ϕ*_(*u*; *Y* ). The invertibility requirement ensures that we can both generate samples (forward pass) and evaluate densities (inverse pass). Complex distributions can be learned by composing simple transformations, each of which maintains invertibility and efficient Jacobian computation. This allows the network to learn highly flexible posterior shapes while maintaining computational tractability, a crucial feature when inferring parameters in complex biological models where posteriors may exhibit non-Gaussian features such as multimodality, heavy tails, or curved degeneracies. The resulting approximate posterior density follows from the change of variables formula

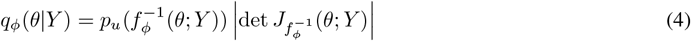

Where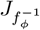denotes the Jacobian of the inverse transformation. Intuitively, the determinant of the Jacobian corrects for the local change in volume caused by the transformation, ensuring the resulting density is properly normalized. To understand this geometrically: when the transformation *f*_*ϕ*_ stretches space in some region, the determinant accounts for this stretching to ensure probability mass is conserved [38]. Modern flow architectures (e.g., coupling flows [40], autoregressive flows [41]) are designed such that this determinant is computationally efficient to calculate— often reducing to a summation or product of scalar terms rather than requiring explicit *d× d* matrix determinant computation. For readers familiar with traditional MCMC-based inference in biological models, normalizing flows offer a complementary approach: rather than exploring the posterior through iterative sampling, they learn an explicit parametric approximation that enables fast posterior evaluation and sampling once trained [21]. This amortization— training once on simulated data, then instantly obtaining posteriors for new observations—makes them particularly attractive for scenarios requiring repeated inference, such as model selection, experimental design, or real-time data analysis.

#### 2.3.3 Training Procedure

NPE training requires a dataset of parameter-data pairs generated from the simulator. The training procedure follows three main steps:

1. **Simulation:** Generate *N* training samples by drawing parameters from the prior *θ*_*i*_ *∼ p*(*θ*) and simulating corresponding data *Y*_*i*_ *∼ p*(*Y* |*θ*_*i*_) using the stochastic model (Figure 2, left).
2. **Optimization:** Train the normalizing flow by maximizing the log-likelihood„ of parameters given their corresponding simulated data,

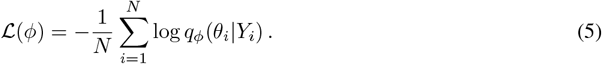

This objective is optimized using stochastic gradient descent (e.g., Adam [42]) with mini-batches of simulated pairs (*θ*_*i*_, *Y*_*i*_). During training, gradients are computed via back-propagation through the flow transformation *f*_*ϕ*_(*u*; *x*), updating the network parameters *ϕ* until convergence. This learns the transformation that maps from a simple base distribution to the posterior distribution conditioned on the data (Figure 2, left).

- **Inference:** For observed data *Y*_obs_, obtain the approximate posterior *q*_*ϕ*_(*θ* |*Y*_obs_) by evaluating the trained network. Generate posterior samples by drawing *u∼ p*_*u*_(*u*) and applying the learned transformation *θ* = *f*_*ϕ*_(*u*; *Y*_obs_) (Figure 2, right).

The training data requirements scale with problem complexity, but it is difficult in general to provide a quantitative recommendation on the number of required simulations to train the model. Typically at least *N* = 10^4^− 10^6^ simulations suffice for moderate-dimensional parameter spaces (*d*≤ 10), though higher-dimensional problems may require larger datasets [43]. Crucially, this procedure is amortized; the computationally expensive training phase is performed only once, after which posterior inference for any new observation is nearly instantaneous. This represents a significant advantage over methods like ABC, which require fresh simulations for each new dataset. We provide quantitative comparisons in Section 3.6.

#### 2.3.4 Implementation Details

All NPE analyses in this work use the `sbi` Python package [44], which provides a well-documented implementation of state-of-the-art simulation-based inference algorithms. We employ neural spline flows [45] as our density estimator architecture. Neural spline flows construct invertible transformations from monotonic rational-quadratic spline segments, providing highly expressive density estimation with analytically tractable Jacobians and stable training dynamics. We use single-round NPE throughout; sequential variants [19, 20] can improve sample efficiency when the prior is much broader than the posterior, but our prior ranges are sufficiently constrained that single-round training provides accurate posteriors without sequential refinement. Additional implementation details can be found in the project GitHub repository^2^.

Prior distributions for each model are specified alongside the corresponding inference results in Section 3. In all cases we use broad uniform priors determined by the physical constraints of the model parameters; for NPE, the prior serves the dual role of defining the training distribution from which simulation parameters are drawn and entering into the posterior distribution via Bayes’ theorem [36, 46].

### 2.4 Data Representations and CNN Architecture

The barrier assay experimental design naturally lends itself to a dimensional reduction from two-dimensional spatial data to one-dimensional measurements. Since the initial macroscopic density is uniform in the vertical (*y*) direction and boundary conditions preserve this symmetry, the resulting agent distribution remains independent of vertical location on average. This motivates quantifying the system by counting the number of agents per column

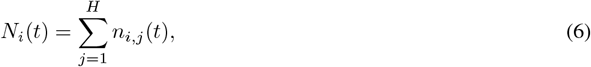

where *n*_*i,j*_(*t*) denotes the integer number of agents at lattice site (*i, j*) at time *t*, and *H* is the number of lattice sites in the vertical direction. This dimensional reduction from 2D spatial data to 1D column counts is standard in cell biology scratch assays [1], as it simplifies the characterization of spatio-temporal spreading while retaining the stochastic fluctuations inherent to the discrete process, as evident in Figure 1(c)–(d). The continuous analogue of the column count *N*_*i*_ under the PDE model is *Hu*(*x*_*i*_, *t*), where *x*_*i*_ denotes the horizontal position of the *i*-th column and *H* is the lattice height. Figure 1(c)–(d) compares the stochastic count data (blue dots) with this PDE solution (red curves), revealing that while the PDE captures mean trends, substantial stochastic fluctuations remain.

While column counts are the standard data representation for barrier assays, this dimensional reduction necessarily discards spatial information—such as clustering patterns, local density fluctuations, and correlations in the vertical direction—that may help constrain parameters in models where processes like proliferation generate spatial structure not captured by one-dimensional projections (see Section 3.1.3). We therefore also develop a pipeline that operates directly on the full two-dimensional spatial data. We treat each lattice configuration from the simulator as a 2D histogram (or count-based image), where each pixel’s value represents the integer number of agents *n*_*i,j*_(*t*) at the corresponding lattice site. To learn informative features directly from this high-dimensional input, we replace the simple feed-forward network with a CNN [47, 48, 49]. The CNN acts as a learnable feature extractor embedded within the NPE pipeline. It uses a series of convolutional and pooling layers to progressively downsample the spatial data into a low-dimensional embedding. This process captures salient spatial features, such as clustering patterns and local agent densities, while preserving translational invariance. The learned embedding is then passed to the remainder of the network to estimate the posterior distribution. This end-to-end training ensures that the features extracted by the CNN are optimized for parameter inference.

Table 2 summarises the key hyperparameters for both the 1D and 2D NPE configurations. A detailed specification of the CNN architecture and full implementation details can be found in the project GitHub repository.

**Table 2:**
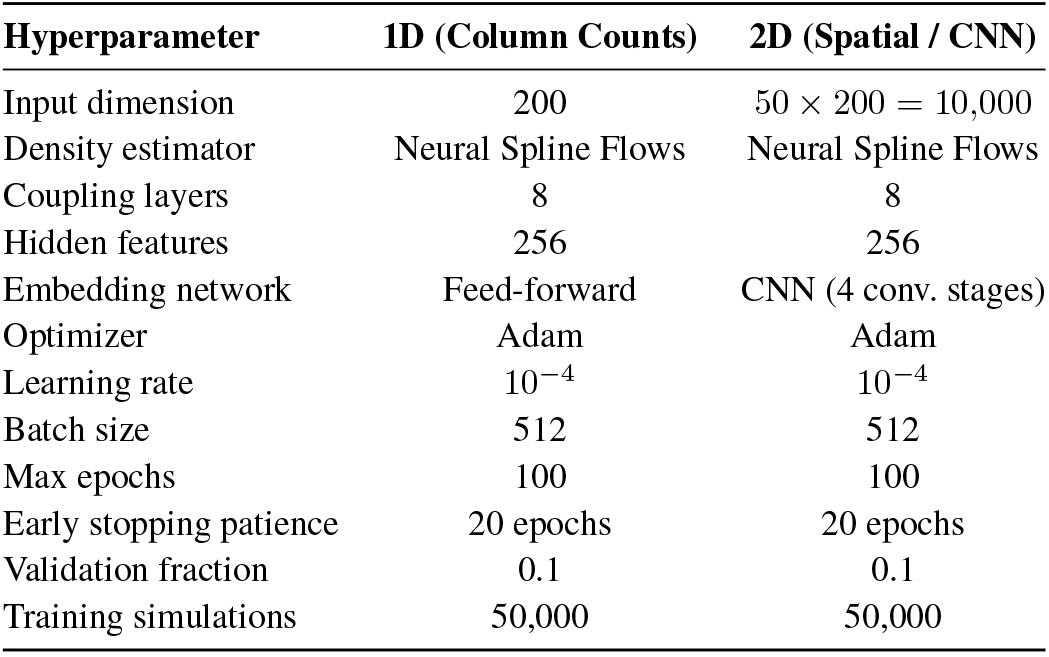
Summary of key hyperparameters for the single-round NPE pipeline in the 1D and 2D configurations. All models were trained on an NVIDIA A100-SXM4-80GB GPU using the sbi library [44] with a PyTorch backend.

| Hyperparameter | 1D (Column Counts) | 2D (Spatial / CNN) |
| --- | --- | --- |
| Input dimension | 200 | $50 \times 200 = 10,000$ |
| Density estimator | Neural Spline Flows | Neural Spline Flows |
| Coupling layers | 8 | 8 |
| Hidden features | 256 | 256 |
| Embedding network | Feed-forward | CNN (4 conv. stages) |
| Optimizer | Adam | Adam |
| Learning rate | $10^{-4}$ | $10^{-4}$ |
| Batch size | 512 | 512 |
| Max epochs | 100 | 100 |
| Early stopping patience | 20 epochs | 20 epochs |
| Validation fraction | 0.1 | 0.1 |
| Training simulations | 50,000 | 50,000 |

## 3 Results

We present inference results for four progressively complex random walk models. We begin with the original isotropic model (Section 3.1), which serves as a baseline comparison with classical inference methods. We then extend the analysis to models incorporating directional bias (Section 3.2), proliferation (Section 3.3), and their combination (Section 3.4). For each model, we report NPE posteriors from both 1D column count and 2D spatial data. Section 3.5 validates all posteriors using simulation-based calibration diagnostics and Section 3.6 compares computational costs.

### 3.1 Original Model

For the original isotropic model, we infer two parameters: the initial occupation probability *U* (cell seeding density) and the motility probability *P*, where the effective diffusion coefficient is *D* = *P/*4. In all cases, inference is performed using a single snapshot of simulated data at *t* = 100 time steps. We use ground truth values *U* = 0.3 and *D* = 0.175.

#### 3.1.1 Classical Inference Baseline

Figure 3a shows the posterior obtained via ABC. This method operates directly on the stochastic model but relies on one-dimensional summary statistics (column counts), leading to a notably broad posterior. This diffuseness reflects both the inherent stochasticity of the system and the information loss from summarizing the data, showcasing the difficulty in precisely constraining parameters even with a high computational budget. In contrast, Figure 3b shows the result of repeating the same ABC rejection algorithm using a deterministic PDE surrogate in place of the stochastic simulator. Because the surrogate is noise-free, parameter combinations that are even slightly inconsistent with the data are rejected, yielding a much tighter posterior without requiring a noise model or estimating *σ*. However, this apparent precision is entirely conditional on the fidelity of the surrogate; any mismatch between the surrogate and the true stochastic process introduces a systematic bias not reflected in the narrow posterior. Finally, Figure 3c shows the posterior sampled using MCMC with the surrogate model and an assumed Gaussian noise model. This provides a more flexible characterization of uncertainty. Crucially, this method requires inferring an additional nuisance parameter, *σ*, which quantifies the noise variance. Figure 3c shows the joint posterior over the model parameters and this additional noise term. While comprehensive, this result is still limited by the choice of surrogate and the specified noise model.

**Figure 3.**
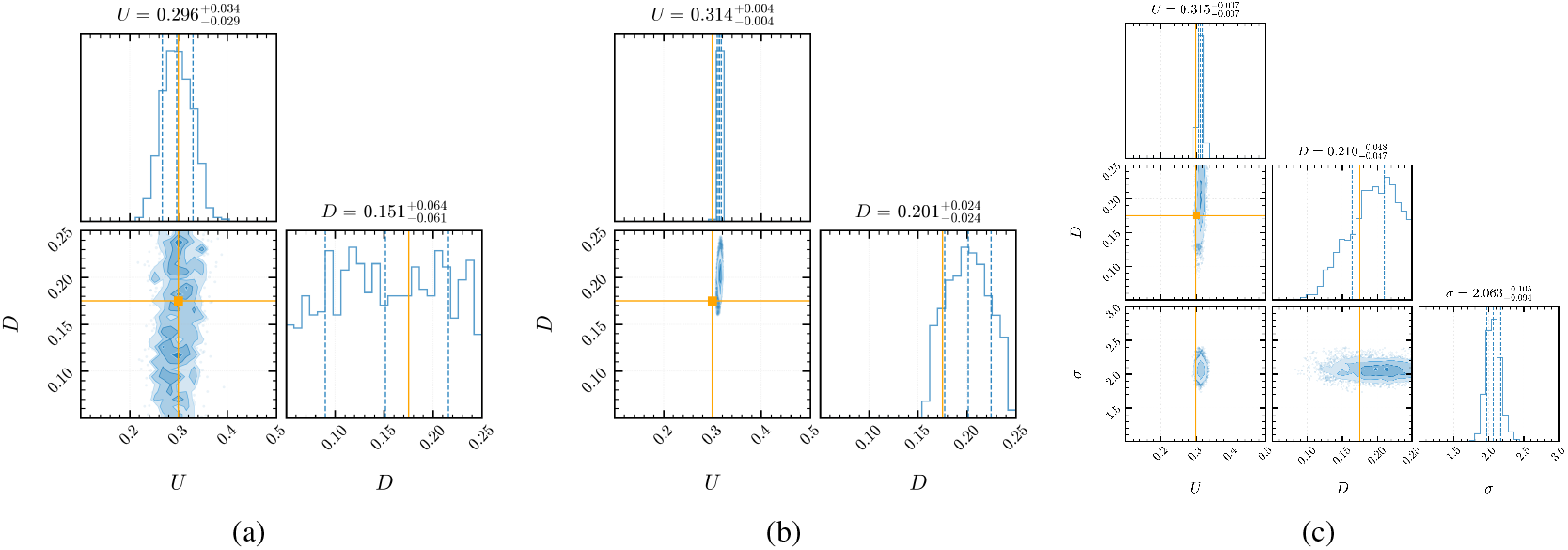
Comparison of posterior distributions obtained using classical inference methods for a scratch assay model. (a) Approximate posterior from ABC applied directly to the stochastic model; computational constraints and information loss lead to a diffuse posterior for *D*. (b) Approximate posterior from ABC using a deterministic PDE surrogate in place of the stochastic simulator; the noise-free surrogate yields a tighter posterior but is conditional on the surrogate’s fidelity. (c) Posterior distribution from MCMC using the surrogate model with an assumed Gaussian noise model; this method jointly infers the model parameters alongside an additional noise variance parameter, *σ*. In all panels, the diagonal shows the marginalized one-dimensional posteriors for each parameter and the off-diagonal shows the joint two-dimensional posterior. The subtitles above each one-dimensional subplot indicate the posterior median and the central 68% credible interval. The vertical dashed lines mark the 16th, 50th, and 84th percentiles, and the orange lines indicate the true parameter values. No true value is shown for *σ* in panel (c) because it is a nuisance parameter introduced by the surrogate modelling approach; the true data-generating process is the stochastic simulator, which has no corresponding noise variance parameter.

Collectively, these results highlight the central trade-off: direct methods like ABC are computationally demanding and often imprecise, while surrogate-based methods are faster but introduce hard-to-quantify systematic biases that cannot be reduced by collecting more data.

#### 3.1.2 NPE with 1D Summary Statistics

We demonstrate NPE’s effectiveness by applying it to the same synthetic barrier assay data used in Section 3.1.1. To ensure direct comparison with the classical methods, we first implement NPE using identical 1D column count summary statistics *N*_*i*_, representing the number of agents in each vertical column of the lattice. Following the protocol of S&P2025, we generate synthetic observations from a stochastic random walk model with true parameters *P* = 0.7 and *U* = 0.3, corresponding to diffusivity *D ≈ P/*4 = 0.175. The resulting data consists of a 201-dimensional vector of column counts, corresponding to the 201 discrete lattice columns from *x* = −100 to *x* = +100. We perform inference directly on the stochastic model parameters *P* and *U*, then transform to *D* via the relationship *D* = *P/*4 for comparison with PDE-based approaches. NPE recovers a posterior distribution with median *U* = 0.293 *±* 0.008 and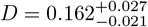 (Figure 4a). The results demonstrate excellent agreement with the true parameter values (*U* = 0.3, *D* = 0.175).

**Figure 4.**
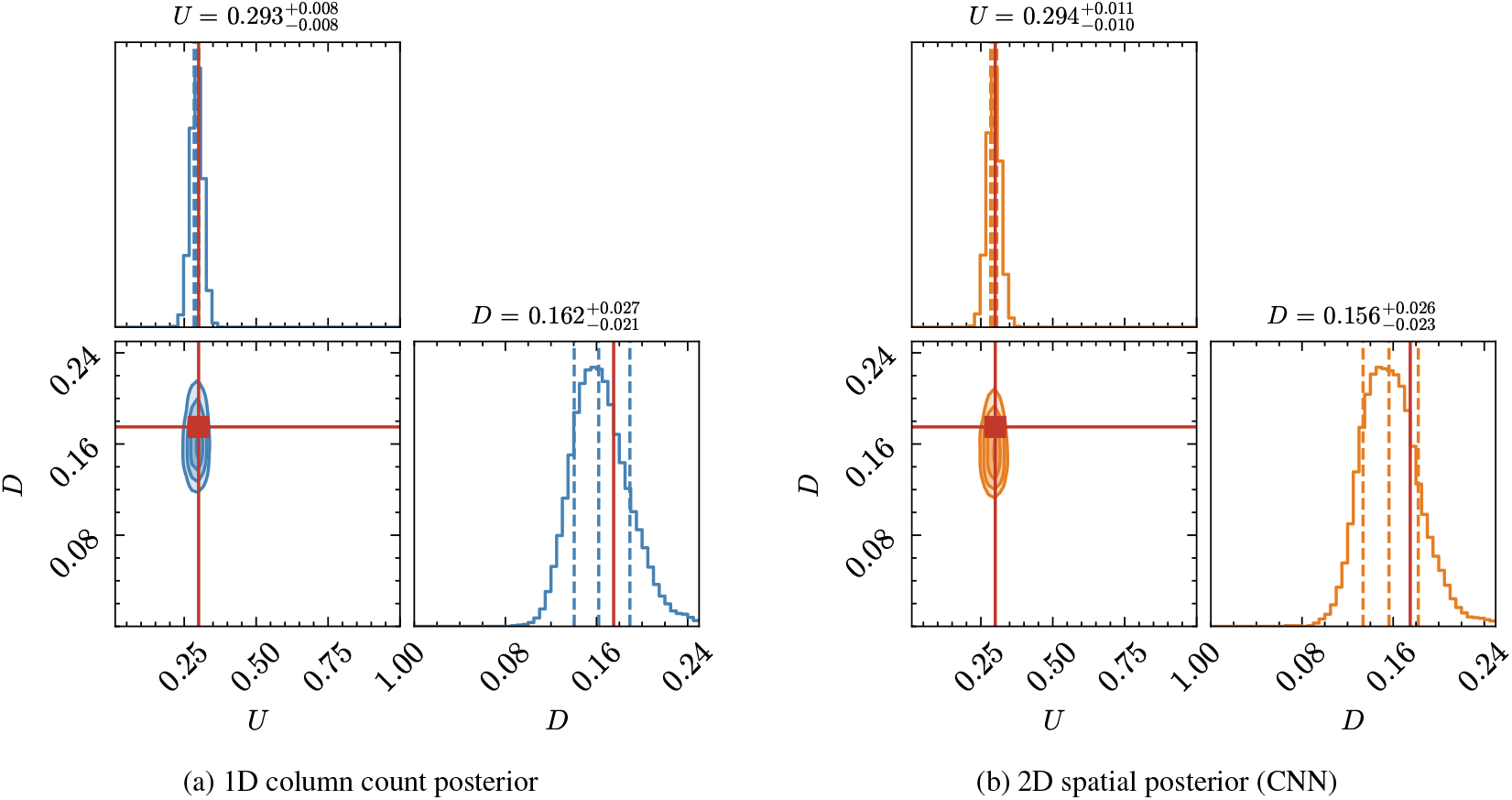
Posterior distributions for the original model with true values *U* = 0.3, *D* = 0.175. (a) NPE posterior from 1D column count data yields *U* = 0.293 *±* 0.008 and 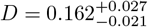 (medians with 68% credible intervals). (b) NPE posterior from 2D spatial data using CNN embedding yields 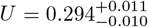 and 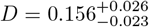 (medians with 68% credible intervals). Both approaches recover comparable posteriors, demonstrating that the CNN automatically extracts features from the raw spatial data that are as informative as the curated 1D summary statistics.

This approach yields several key advantages over classical methods. The NPE posterior shows substantially improved precision relative to ABC, especially for the diffusion parameter *D*, all without the need for manual tolerance tuning. Because NPE learns directly from the stochastic simulator, it avoids the systematic bias of mean-field approximations and produces a posterior that is not artificially narrow. The resulting credible intervals accurately reflect the natural stochastic variance of the data, a feature often lost when using deterministic surrogate models, cf. Figure 3. Once the initial network training is complete, posterior samples for any new observation can be obtained almost instantly.

Posterior predictive checks confirm that the inferred model reproduces the observed data, with 94.8% of observations falling within the 95% prediction intervals. Unlike surrogate-based approaches, the prediction intervals are physically consistent—respecting non-negativity and naturally capturing heteroscedastic uncertainty (Appendix A.1).

#### 3.1.3 NPE with 2D Spatial Data

While column counts provide a natural summary of the spreading process, they necessarily discard information about the vertical distribution of agents. In principle, the spatial correlation structure within the 2D configuration contains additional information about the underlying parameters. For instance, clustering patterns, local density fluctuations, and spatial correlations may help distinguish between different combinations of *P* and *U* that produce similar column counts, cf. parameter identifiability^3^. For example, models incorporating agent-agent interactions, complex boundary conditions, spatially heterogeneous environments, or multi-species dynamics would generate spatial patterns that cannot be adequately summarized by simple column counts.

The experimental setup remains identical to the 1D case, using the same synthetic observation generated with true parameters *P* = 0.7 (*D* = *P*/4 = 0.175) and *U* = 0.3. However, instead of using the 201-dimensional vector of column counts as input, the network now receives the full 50*×* 200 binary image representing the agent locations on the lattice. We use the same broad uniform priors for *P* and *U* as described in Section 2.3.

The CNN-based approach recovers a posterior with medians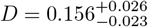 and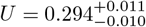(Figure 4b). As summarised in Table 3, leveraging the full spatial data yields a posterior with a precision comparable to that obtained using curated 1D summary statistics, cf. Section 3.1.2.

**Table 3:** Comparison of inference results across methods and data representations. All values show posterior medians with 68% credible intervals (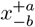 ±denotes symmetric intervals where the upper and lower bounds are equal to the reported precision); classical method posteriors are extracted from the corner plots in Figure 3. The MCMC method additionally infers a noise variance nuisance parameter *σ* (not shown). The surrogate-based methods (ABC with surrogate, MCMC) yield biased estimates of *D*, reflecting systematic error from the mean-field approximation.

| Method | Data Type | Posterior for $U$ | Posterior for $D$ |
| --- | --- | --- | --- |
| True value | — | 0.300 | 0.175 |
| ABC | 1D Column counts | $0.296^{+0.034}_{-0.029}$ | $0.151^{+0.064}_{-0.061}$ |
| Surrogate + ABC | 1D Column counts | $0.314^{+0.004}_{-0.004}$ | $0.201^{+0.024}_{-0.024}$ |
| Surrogate + MCMC | 1D Column counts | $0.315^{+0.007}_{-0.007}$ | $0.210^{+0.048}_{-0.047}$ |
| NPE | 1D Column counts | $0.293 \pm 0.008$ | $0.162^{+0.027}_{-0.021}$ |
| NPE + CNN | 2D Spatial data | $0.294^{+0.011}_{-0.010}$ | $0.156^{+0.026}_{-0.023}$ |

This result demonstrates that the CNN can automatically learn features from the raw data that are as informative as the carefully chosen summary statistics. Importantly, the CNN adds negligible overhead to the network training phase compared to the feed-forward embedding used for 1D data (Section 3.6), so the primary cost of adopting the 2D pipeline is conceptual rather than computational. For more complex models where optimal summary statistics are unknown, this direct-from-pixels approach may be advantageous for accurate inference.

Posterior predictive checks for the 2D model similarly confirm well-calibrated predictions that capture the spatial characteristics of the agent-based simulation (Appendix A.2).

Biologically, *U* represents the initial cell seeding density and *D* the effective cell motility. For this isotropic model, the comparable precision of 1D and 2D posteriors is expected: column counts capture the symmetric spreading dynamics that encode both parameters. The advantage of spatial analysis emerges for models with directional structure, as we demonstrate next (Section 3.2).

### 3.2 Model A — Directional Bias

Model A extends the isotropic random walk by introducing a directional bias parameter *ρ∈* [ −1, 1], which controls the probability of stepping in the positive *x*-direction relative to the negative *x*-direction (see Section 2.1). This model has three free parameters: the initial occupation probability *U*, the total motility probability *P*, and the bias *ρ*, with true values *U* = 0.5, *P* = 0.7, *ρ* = 0.5.

Using 1D column count data, NPE recovers posteriors *U* = 0.498 *±* 0.011, *P* = 0.631 *±* 0.176, and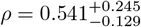(Figure 5a). The initial density *U* is tightly constrained, while *P* and *ρ*exhibit broader posteriors with a strong anti-correlation (*r* = − 0.962). This degeneracy is physics-driven: the macroscopic drift velocity *v* = *Pρ/*2 is well-constrained by the data, but individual values of *P* and *ρ* that yield the same drift are interchangeable when viewed through 1D column counts alone. That is, the composite quantity *v* is identifiable, but *P* and *ρ* are individually weakly identifiable from 1D data.

**Figure 5.**
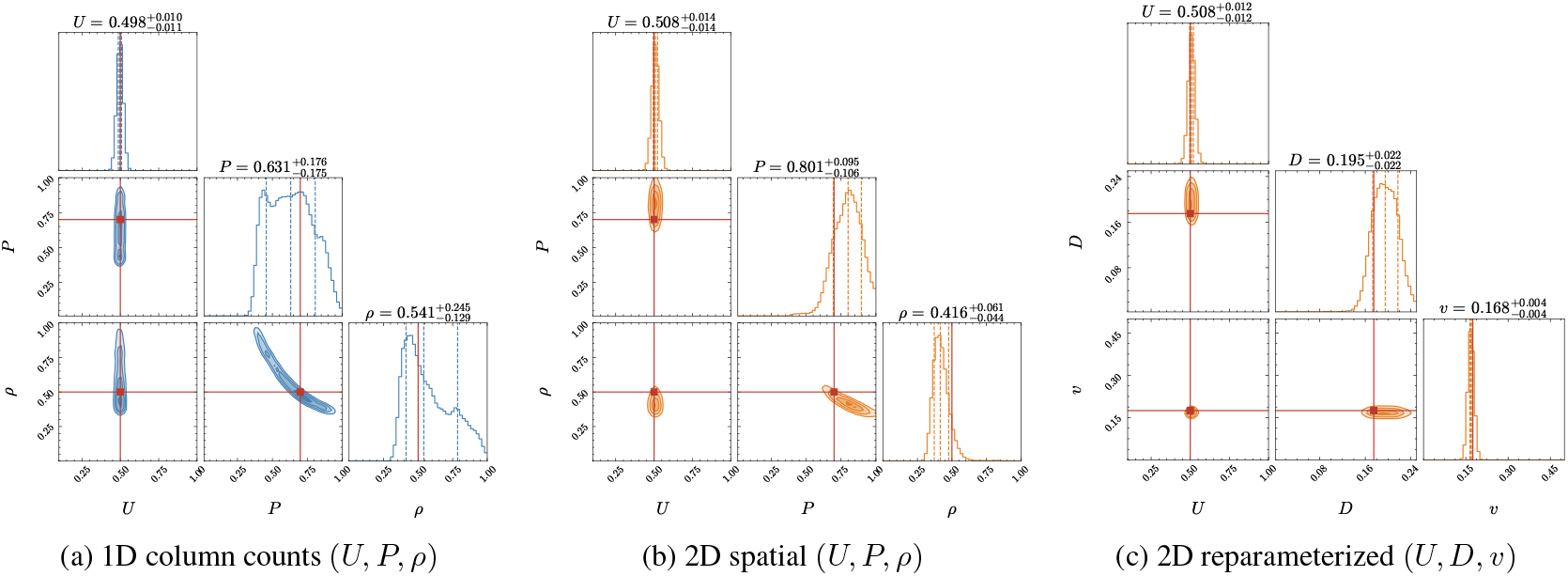
Posterior distributions for Model A (directional bias) with true values *U* = 0.5, *P* = 0.7, *ρ* = 0.5. (a, b) Both 1D and 2D representations recover *U* precisely but exhibit a strong *P* –*ρ* anti-correlation reflecting the drift velocity degeneracy *v* = *Pρ/*2. (c) Reparameterizing to continuum parameters (*U, D, v*), where *D* = *P/*4 and *v* = *Pρ/*2,transforms the banana-shaped degeneracy into a near-Gaussian posterior, with *v* constrained to ∼3.4% relative error.

Using full 2D spatial data with a CNN embedding network, NPE recovers *U* = 0.508 *±* 0.030, 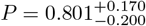 and 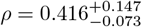 (Figure 5b). The *P* –*ρ* anti-correlation persists (*r* = −0.924), despite the CNN having access to the spatial asymmetries that column counts collapse by summing across rows. This confirms that the degeneracy reflects an intrinsic limitation of single-timepoint inference rather than an information loss artefact of the 1D summary. The 2D medians for *P* and *ρ* lie in a different region of the degenerate manifold compared to the 1D case, reflecting the sensitivity of the posterior mode to the data representation when parameters are weakly identifiable. Biologically, *ρ* quantifies the directional migration preference (e.g. chemotactic bias); the strong *P* –*ρ* coupling means the net directed migration rate is the identifiable and biologically relevant quantity for predicting cell assay closure direction and speed.

Since the drift velocity *v* = *Pρ/*2 is the identifiable quantity, a natural strategy is to reparameterize the NPE to infer continuum parameters (*U, D, v*) directly, where *D* = *P/*4 is the diffusivity and *v* = *Pρ/*2 is the drift velocity. This transformation reuses the same 50,000 training simulations—only the parameter columns are transformed; the observations are unchanged. In the reparameterized space the NPE recovers *U* = 0.508*±* 0.014, *D* = 0.195*±* 0.021 (true 0.175), and *v* = 0.169*±* 0.006 (true 0.175), with *v* constrained to∼ 3.4% relative error (Figure 5c). The banana-shaped *P* –*ρ* posterior (Figure 5b) becomes a near-Gaussian *D*–*v* posterior (Figure 5c), which normalizing flows can represent faithfully with fewer transforms. This simpler posterior geometry also yields substantially improved simulation-based calibration (SBC) diagnostics, as we discuss in Section 3.5. More broadly, when parameter degeneracies are understood *a priori*, reparameterizing to physically identifiable quantities offers a simple and effective route to improved flow calibration.

### 3.3 Model B — Proliferation

Model B incorporates cell proliferation via a Fisher–Kolmogorov mechanism: at each time step, an agent may divide into a daughter cell at an adjacent site with probability *R*. Both motility and proliferation events are subject to an exclusion constraint that prevents multiple occupancy at any lattice site. This model has three free parameters: the initial occupation probability *U*, the motility probability *P*, and the proliferation rate *R*, with true values *U* = 0.5, *P* = 0.7, and *R* = 0.05.

NPE with 1D column counts recovers 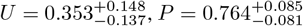, and *R* = 0.052 *±* 0.006 (Figure 6a). While *P* and *R* are tightly constrained, *UR*: higher exhibits a moderately broad posterior reflecting a partial degeneracy with initial occupancy can mimic the effect of increased proliferation when only a single time point is observed.

**Figure 6.**
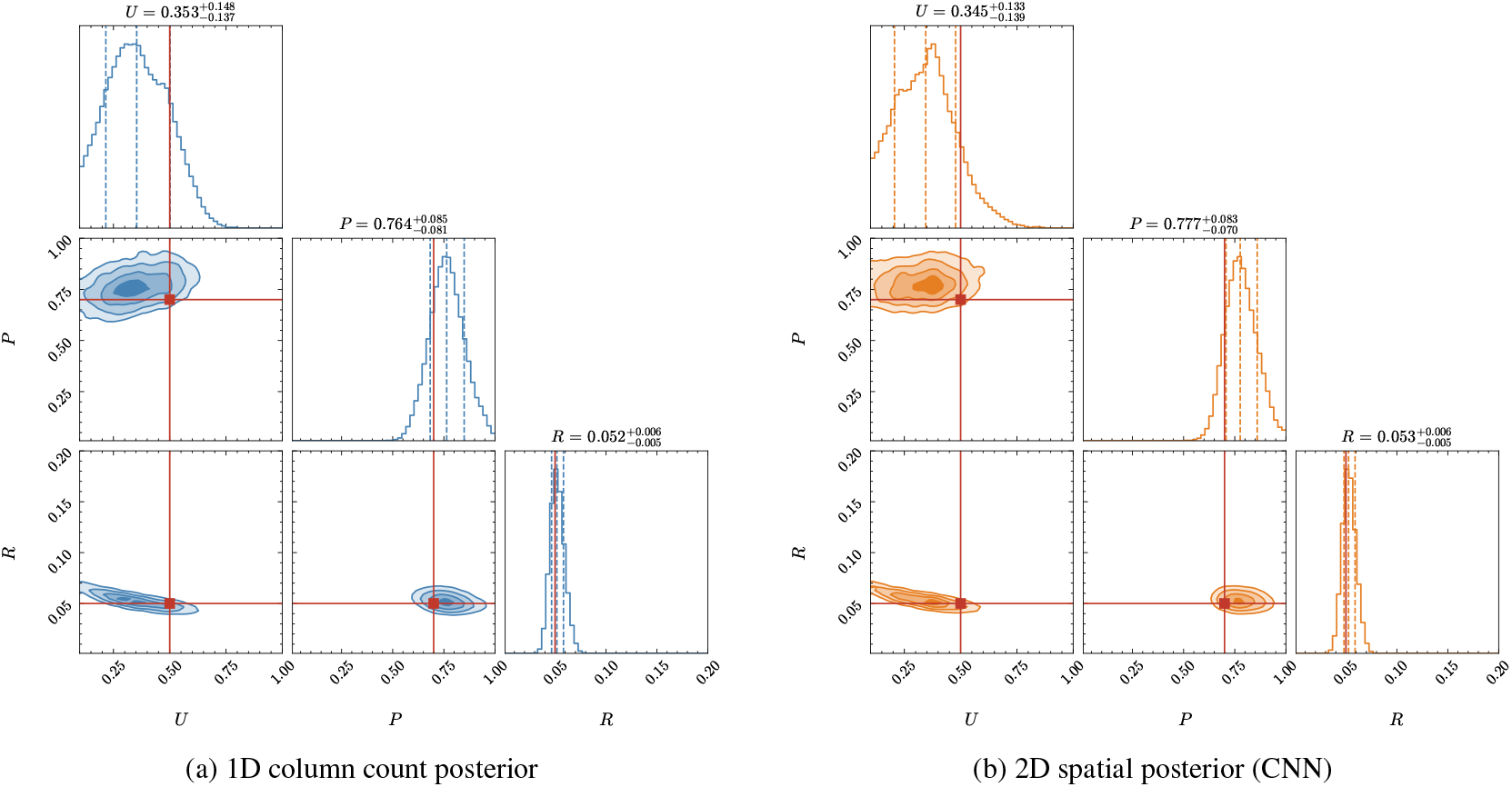
Posterior distributions for Model B (proliferation) with true values *U* = 0.5, *P* = 0.7, *R* = 0.05. Both 1D and 2D representations recover *P* and *R* precisely, while *U* exhibits a partial degeneracy with *R* reflecting the interchangeability of initial density and proliferation for inference using a single time point.

With 2D spatial data, NPE recovers 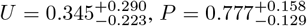, and 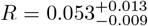 (Figure 6b). The 1D posteriors are at least as tight as the 2D posteriors for all three parameters, suggesting that column counts already capture the information relevant to this proliferation model and that the higher-dimensional 2D observation space may slightly impede flow training. Biologically, the ability to jointly infer *U, P*, and *R* from a single experimental snapshot is valuable: in wound healing assays the seeding density may not be precisely controlled, and NPE can recover it alongside the motility and proliferation parameters without additional observations.

### 3.4 Model C — Combined Bias and Proliferation

Model C combines directional bias and proliferation, representing the most biologically realistic scenario in our suite. Unlike Models A and B, which each introduce one additional mechanism, Model C incorporates both bias and proliferation simultaneously, yielding a four-parameter inference problem: the initial occupation probability *U*, the motility probability *P*, the directional bias *ρ*, and the proliferation rate *R*, with true values *U* = 0.5, *P* = 0.7, *ρ* = 0.5, *R* = 0.05. Although a mean-field PDE can in principle be written for this combination of mechanisms, the interplay of bias, proliferation, and exclusion effects means that the surrogate approximation is no longer well validated, and the additional requirement of specifying an appropriate noise model further compounds the modelling uncertainty. This model therefore represents the strongest case for simulation-based inference, which bypasses surrogate approximations entirely.

NPE with 1D column counts recovers 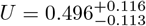, *P* = 0.671 *±* 0.073,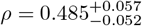 and *R* = 0.052 *±* 0.003(Figure 7a). Despite the four-parameter space,*R* is recovered with∼6% relative uncertainty, and the *P* –*ρ* anti-correlation mirrors the degeneracy observed in Model A.

**Figure 7.**
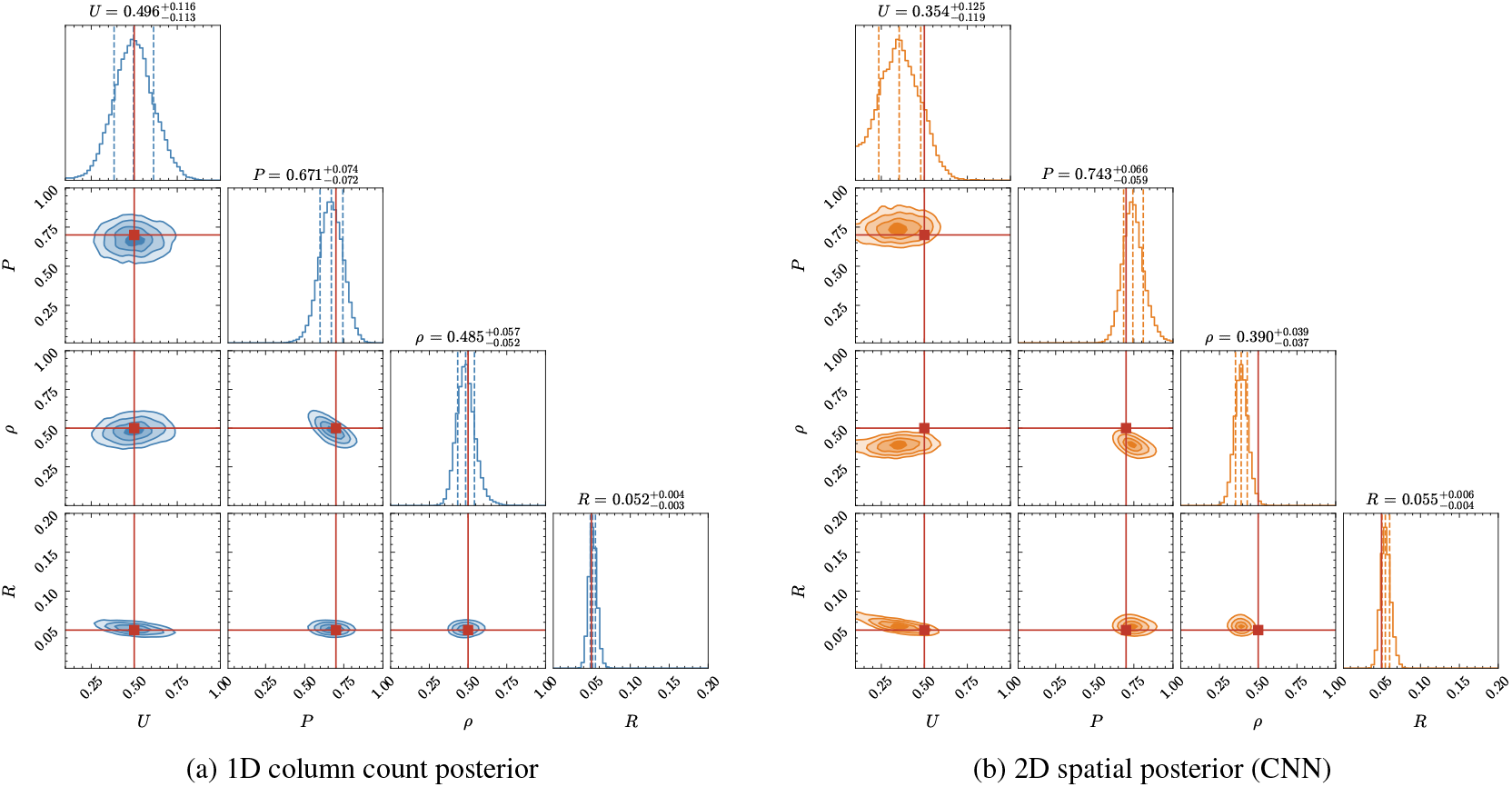
Posterior distributions for Model C (bias + proliferation) with true values *U* = 0.5, *P* = 0.7, *ρ* = 0.5, *R* = 0.05. The *P* –*ρ* anti-correlation from Model A persists, and *R* remains tightly constrained despite the expanded parameter space. The 2D representation yields tighter posteriors for *P* and *ρ*.

With 2D spatial data, NPE recovers 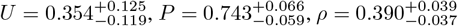, and 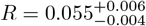 (Figure 7b). The 2D posteriors are tighter for *P* and *ρ*, suggesting that spatial information provides additional constraining power for these parameters, while *U* and *R* are comparably constrained by column counts alone. Biologically, these four parameters represent the key processes governing wound healing dynamics: initial cell density, motility, directional migration, and cell division, all inferred simultaneously from a single experimental snapshot.

### 3.5 SBI Diagnostics

Following the diagnostic protocol recommended by [21] and [43], we validate all trained posteriors using three complementary diagnostics: the Kolmogorov–Smirnov (KS) test for marginal calibration, the classifier two-sample test (C2ST) for posterior quality, and Tests of Accuracy with Random Points (TARP) for coverage calibration.

Table 4 summarises the diagnostic results for both 1D and 2D posteriors across all models. The C2ST scores are consistently close to 0.5 across all models, parameters, and data representations, indicating that the learned posteriors are nearly indistinguishable from the true posteriors and that the density estimators have been well-trained. The TARP area-under-the-curve (AUC) metric passes for the Original model and Model A in both 1D and 2D (AUC values near zero), while Models B and C show larger deviations in the 1D setting (AUC = − 0.44 and − 0.54, respectively), indicating moderate miscalibration for the proliferation models.

**Table 4:** Diagnostic summary for 1D and 2D NPE posteriors across all models. KS *p*-values marked with ^*\**^ indicate failure (*p* < 0.05). C2ST scores near 0.5 indicate well-learned posteriors. TARP AUC values near zero indicate good calibration.

| Model | Param. | 1D NPE |  |  | 2D NPE (CNN) |  |  |
| --- | --- | --- | --- | --- | --- | --- | --- |
| | | KS $p$ | C2ST | AUC | KS $p$ | C2ST | AUC |
| Original | $U$ | 0.96 | 0.59 | 0.05 | 0.0001* | 0.58 | −0.41 |
| | $P$ | 0.0001* | 0.58 | | 0.02* | 0.59 | |
| A | $U$ | 0.0005* | 0.57 | −0.05 | 0.0000* | 0.64 | 0.08 |
| | $P$ | 0.07 | 0.56 | | 0.0000* | 0.56 | |
| | $\rho$ | 0.15 | 0.57 | | 0.02* | 0.58 | |
| B | $U$ | 0.009* | 0.58 | −0.44 | 0.11 | 0.57 | −0.36 |
| | $P$ | 0.0000* | 0.59 | | 0.41 | 0.57 | |
| | $R$ | 0.0000* | 0.57 | | 0.009* | 0.59 | |
| C | $U$ | 0.03* | 0.56 | −0.54 | 0.004* | 0.58 | −0.12 |
| | $P$ | 0.02* | 0.56 | | 0.07 | 0.58 | |
| | $\rho$ | 0.36 | 0.59 | | 0.0000* | 0.58 | |
| | $R$ | 0.0000* | 0.56 | | 0.22 | 0.58 | |

For the 1D posteriors, the KS test results are mixed across models. In the Original model, *U* passes comfortably (*p* = 0.96) while *P* fails (*p* = 0.0001). In Model A, *U* fails (*p* = 0.0005), *P* passes marginally (*p* = 0.07), and *ρ* passes (*p* = 0.15). Model B shows KS failures for all three parameters (*U, P*, and *R*), and Model C fails for *U, P*, and *R* while *ρ* passes (*p* = 0.36). The KS failures are concentrated in the proliferation models (B and C), where the SBC rank empirical cumulative distribution functions (ECDFs; Figure 8c–d) show systematic deviations characteristic of mild overconfidence: the learned posteriors are precise but slightly too narrow, so that the true value falls in the tails more often than expected. This likely reflects the nonlinear and relatively flat mapping from parameters to observable column counts in the proliferation models: modest changes in *R* or *U* produce only subtle differences in the data, making it difficult for the normalizing flow to accurately learn the posterior width.

**Figure 8.**
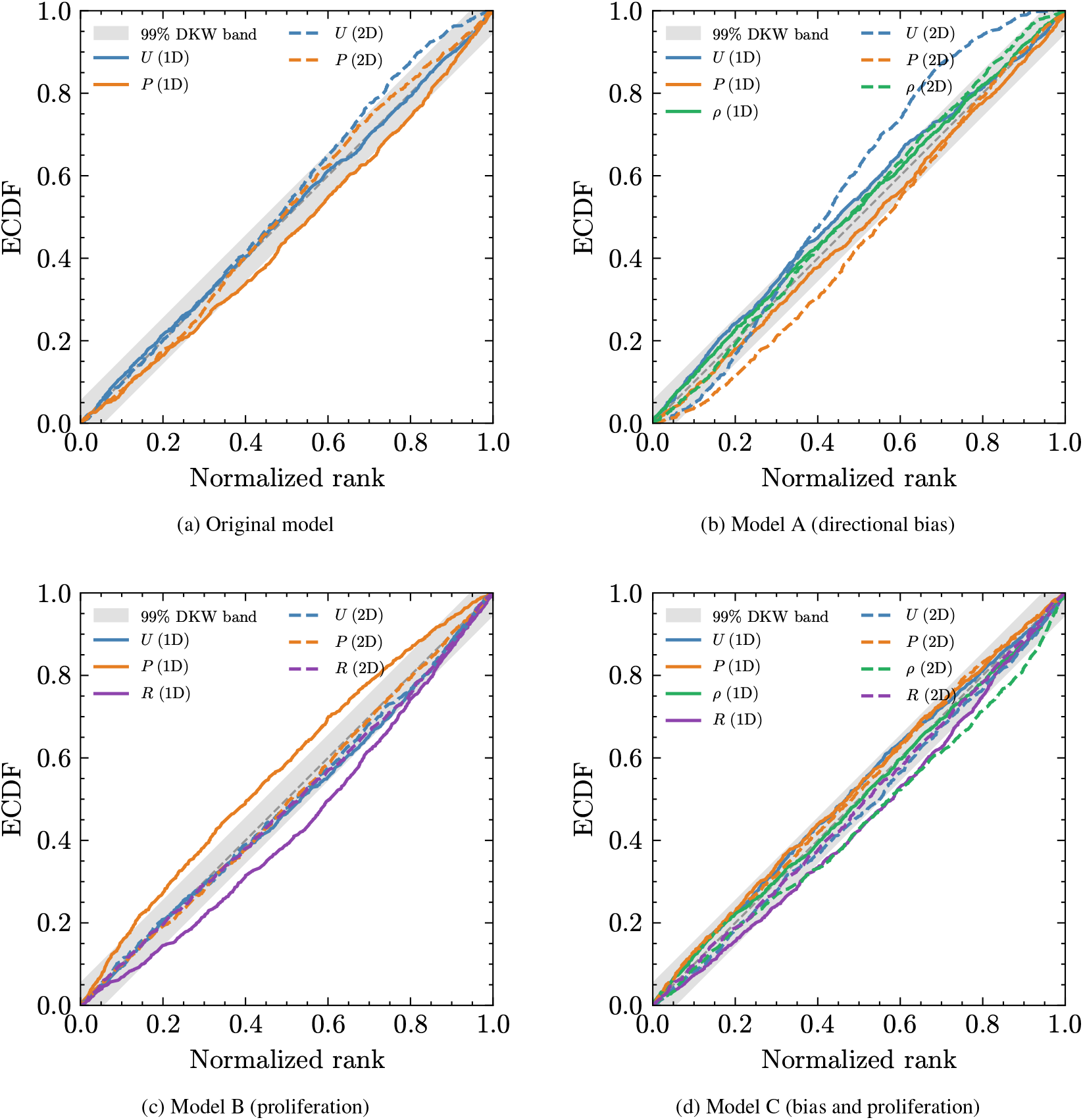
SBC rank ECDF plots for all four models. Solid lines show 1D NPE posteriors; dashed lines show 2D NPE (CNN) posteriors. A uniform rank distribution (diagonal within the shaded 99% DKW confidence bands) indicates well-calibrated posteriors; an ECDF bowing below the diagonal indicates mild overconfidence. (a) Original model: *U* is well-calibrated in 1D while *P* deviates; 2D posteriors show deviations for both parameters. (b) Model A: *U* fails in both representations; *P* and *ρ* pass in 1D, while all three parameters fail in 2D. (c) Model B: all three parameters (*U, P, R*) fail the KS test in 1D; *R* shows marginal deviation in 2D. (d) Model C: the four-parameter space shows KS failures for *U, P*, and *R* in 1D (with *ρ* passing), and for *U* and *ρ* in 2D.

The 2D posteriors exhibit broadly similar calibration patterns (dashed lines in Figure 8), with C2ST scores remaining near 0.5 throughout. The KS test is somewhat more sensitive in the 2D setting: the Original model shows failures for both *U* and *P* ; Model A fails for all three parameters (*U, P*, and *ρ*); Model B shows a marginal failure for *R*; and Model C fails for *U* and *ρ* while *P* and *R* pass. These failures likely reflect the challenge of learning accurate posterior widths in the higher-dimensional observation space. For Model A, where *P* and *ρ* are strongly degenerate, the KS failures may additionally reflect the CNN concentrating posterior mass on a different region of the degenerate manifold than the 1D embedding, as evidenced by the shifted marginal medians (Section 3.2).

This tendency to underestimate posterior variance is a recognised challenge across simulation-based inference methods more broadly [50], though amortized approaches such as NPE tend to be more conservative and more simulation-efficient than their non-amortized counterparts. The KS test is highly sensitive to subtle deviations in the tails of the rank distribution, which may not affect practical inference quality; by contrast, the C2ST diagnostic passes consistently across all models and parameters, providing stronger evidence that the learned posteriors are suitable for practical use. Increasing the training budget or employing sequential refinement strategies could improve KS calibration further, but the miscalibration observed here is mild, with curves remaining close to the Dvoretzky–Kiefer–Wolfowitz (DKW) confidence bands. We additionally verify that these results are robust to the choice of random seed in Appendix B.

When parameter degeneracies are understood *a priori*, reparameterizing to identifiable quantities can substantially improve calibration. For Model A, reparameterizing from lattice parameters (*U, P, ρ*) to continuum parameters (*U, D, v*) (Section 3.2) yields markedly better diagnostics (Figure 9): *D* now passes the KS test (*p* = 0.60, versus *p <* 0.001 for *P* in the baseline), the *U* C2ST score improves from 0.64 to 0.60, the TARP AUC halves from 0.084 to 0.035, and all six C2ST tests pass (versus five of six in the baseline; see also Table 4). The improvement arises because the reparameterization transforms the curved *P* –*ρ* banana-shaped posterior into a near-Gaussian *D*–*v* posterior that the normalizing flow represents more faithfully.

**Figure 9.**
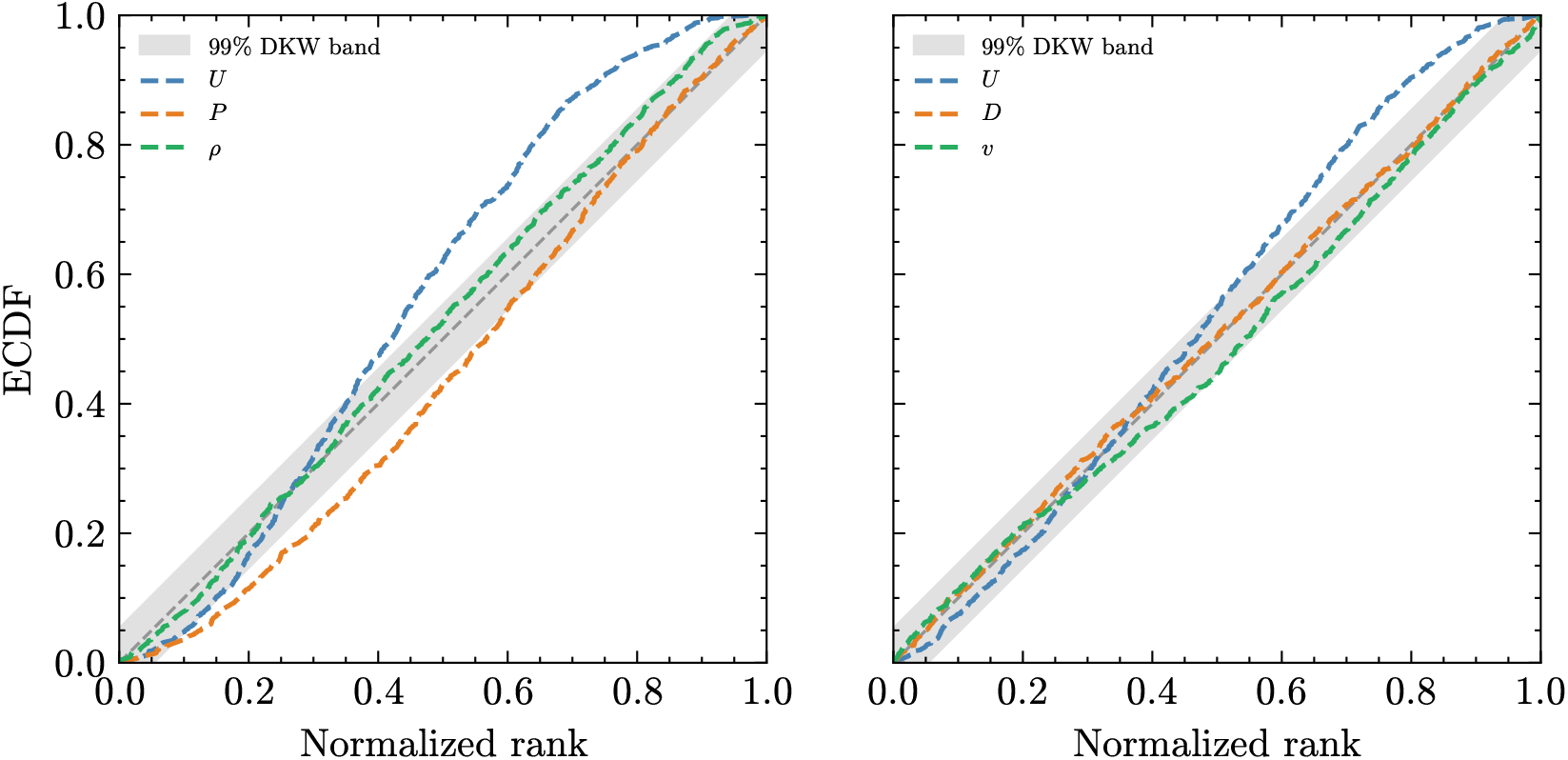
SBC rank ECDFs for Model A (2D): baseline parameterization (*U, P, ρ*) (left) versus reparameterized (*U, D, v*) (right). Shaded bands show the 95% DKW confidence envelope. The left panel reproduces the 2D (dashed) curves from Figure 8(b) for direct comparison with the reparameterized result.

Table 5 provides a comprehensive summary of NPE inference results across all four models and both data representations.

**Table 5:**
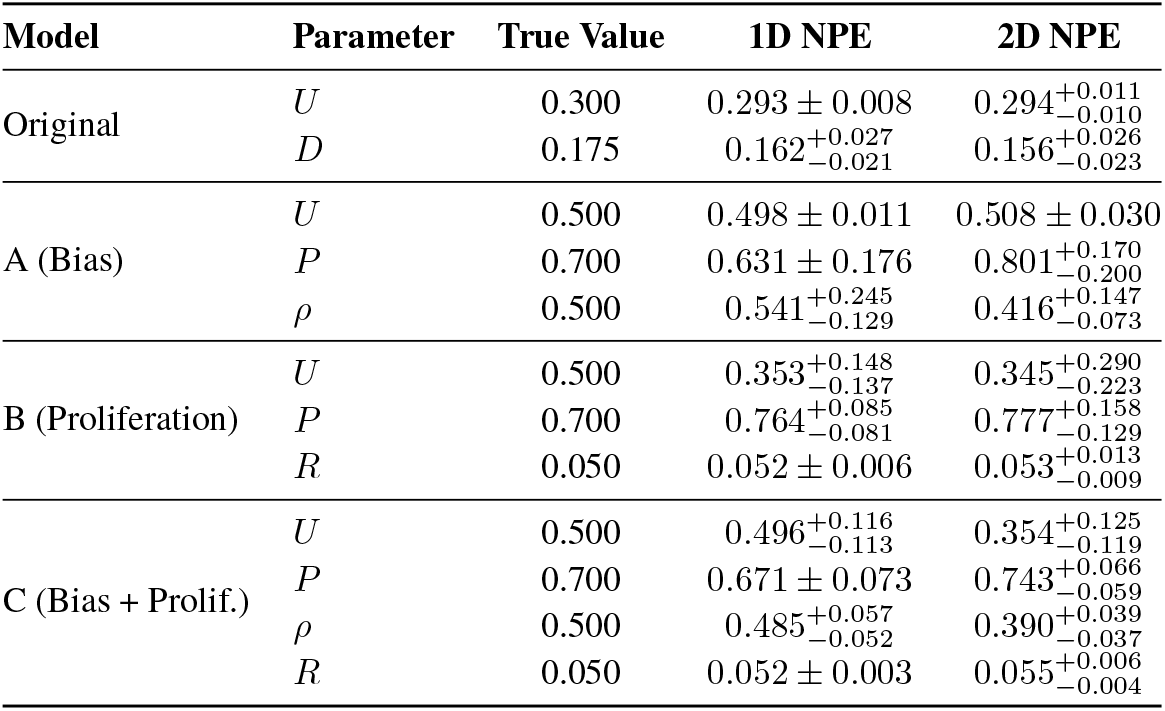
Comprehensive NPE inference results across all models. Values show posterior medians with 68% credible intervals (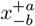; denotes symmetric intervals). For the original model, *D* = *P/*4 is reported for comparison with classical methods. All models use broad uniform priors on the unit interval for probability parameters and *R ∼ U* (0.001, 0.2) for the proliferation rate.

| Model | Parameter | True Value | 1D NPE | 2D NPE |
| --- | --- | --- | --- | --- |
| Original | $U$ | 0.300 | $0.293 \pm 0.008$ | $0.294^{+0.011}_{-0.010}$ |
| | $D$ | 0.175 | $0.162^{+0.027}_{-0.021}$ | $0.156^{+0.026}_{-0.023}$ |
| A (Bias) | $U$ | 0.500 | $0.498 \pm 0.011$ | $0.508 \pm 0.030$ |
| | $P$ | 0.700 | $0.631 \pm 0.176$ | $0.801^{+0.170}_{-0.200}$ |
| | $\rho$ | 0.500 | $0.541^{+0.245}_{-0.129}$ | $0.416^{+0.147}_{-0.073}$ |
| B (Proliferation) | $U$ | 0.500 | $0.353^{+0.148}_{-0.137}$ | $0.345^{+0.290}_{-0.223}$ |
| | $P$ | 0.700 | $0.764^{+0.085}_{-0.081}$ | $0.777^{+0.158}_{-0.129}$ |
| | $R$ | 0.050 | $0.052 \pm 0.006$ | $0.053^{+0.013}_{-0.009}$ |
| C (Bias + Prolif.) | $U$ | 0.500 | $0.496^{+0.116}_{-0.113}$ | $0.354^{+0.125}_{-0.119}$ |
| | $P$ | 0.700 | $0.671 \pm 0.073$ | $0.743^{+0.066}_{-0.059}$ |
| | $\rho$ | 0.500 | $0.485^{+0.057}_{-0.052}$ | $0.390^{+0.039}_{-0.037}$ |
| | $R$ | 0.050 | $0.052 \pm 0.003$ | $0.055^{+0.006}_{-0.004}$ |

### 3.6 Computational Performance

To provide a practical comparison of computational costs, Table 6 separates the NPE cost into simulation generation and neural network training. Simulation generation is the dominant expense, accounting for approximately 74% of the total time for the original model and over 96% for Models A–C. This dominance arises because each training simulation involves running the full stochastic agent-based model: the original model generates ∼20 simulations per second, while Models A–C produce only ∼2 per second due to the additional exclusion and interaction logic. In contrast, neural network training is fast ( ∼2–3 minutes on an A100 GPU) and largely independent of model complexity, since it operates on the already-generated data tensors. The choice of 1D column-count versus 2D spatial-grid output has negligible impact on simulation time, as the underlying stochastic simulation is identical; only the output format differs. The CNN-based 2D pipeline also adds negligible overhead to the network training phase: for a fixed training set size, CNN and feed-forward network training times are comparable on GPU hardware. These figures are illustrative of a typical workflow and are not the result of systematic hyper-parameter optimisation. The simulation budget (here 50,000 samples) scales the generation cost proportionally, with an uninvestigated trade-off in posterior accuracy. Similarly, ABC runtime depends on the number of simulations generated (here, 10^5^). Other hyper-parameters (e.g. number of network epochs, batch size, learning rate, width/depth of the embedding network) also have a direct influence on both runtime and accuracy, but were fixed to reasonable defaults rather than tuned for performance.

**Table 6:** Computational runtimes for various inference methods. NPE costs are separated into simulation generation (50,000 training simulations on 16 CPU workers) and neural network training (on an A100 GPU). Simulation time is independent of output format (1D column counts versus 2D spatial grids), as the underlying stochastic simulation is identical. Simulation generation dominates the cost, accounting for ∼74% of the total for the original model and over 96% for Models A–C. Once trained, NPE inference for 5,000 posterior samples takes under one second.

| Method | Model | Sim. Generation (s) | NN Training (s) | Inference (s) |
| --- | --- | --- | --- | --- |
| ABC | Stochastic Simulator | – | – | ~2000 |
| ABC | Surrogate Model | – | – | ~2 |
| MCMC | Surrogate Model | – | – | ~1 |
| NPE | Original | ~515 | ~175 | <1 |
| NPE | Model A | ~4325 | ~135 | <1 |
| NPE | Model B | ~4925 | ~125 | <1 |
| NPE | Model C | ~4855 | ~170 | <1 |

Two distinct computational strategies emerge from this comparison. Surrogate-based approaches are extremely fast because they entirely bypass repeated stochastic simulations, relying instead on precomputed or deterministic approximations of the model. In contrast, the NPE framework demands a substantial one-time computational investment, dominated by simulation generation rather than network training, but then performs posterior inference for new observations almost instantaneously. This separation is practically important: the simulation bottleneck is inherent to the stochastic model and can be mitigated through parallelism or more efficient simulators, whereas the network training component is already fast and scales well on modern GPU hardware. The amortized structure makes NPE particularly attractive for scenarios requiring repeated or real-time inference across many datasets.

## 4 Discussion

### 4.1 Advantages and Limitations of Neural Posterior Estimation

NPE offers a robust and flexible framework for parameter inference, directly addressing several challenges that have limited the practical application of stochastic agent-based models. By learning directly from the full stochastic simulator, it eliminates the need for surrogate models. This direct approach circumvents the systematic bias introduced by mean-field approximations and removes the challenging requirement of specifying explicit noise models that connect deterministic surrogates to stochastic observations. Normalizing flows can also capture complex, multimodal posterior geometries that arise naturally from model non-identifiabilities without requiring restrictive parametric assumptions about posterior shape. Processing raw high-dimensional data through automatic feature extraction avoids the information loss inherent in manually crafted summary statistics while enabling the analysis of complex spatial patterns that would be difficult to characterize a priori. NPE is also amortized: after the initial investment in generating training data and optimizing the neural network, posterior inference for new observations requires under a second of network evaluation rather than the minutes to hours required by simulation-based methods like ABC.

However, NPE also introduces new challenges and limitations that practitioners must carefully consider. The method’s appetite for training data can be substantial, particularly for high-dimensional parameter spaces where the curse of dimensionality becomes prohibitive. Problems with more than 15–20 parameters may require simulation budgets that exceed practical computational limits. NPE performance can also degrade significantly when test observations fall outside the training distribution, making careful prior specification and robust training data generation critical for reliable inference.

The neural network architecture introduces another layer of complexity, as design choices regarding depth, width, and flow type can dramatically impact performance. Unlike classical methods with well-established theoretical foundations, NPE provides limited formal guarantees about convergence or approximation quality. This black-box nature can make diagnosis of poor performance challenging and requires practitioners to develop intuition about network behaviour and training dynamics.

Our model hierarchy illustrates a spectrum from convenience to necessity for simulation-based inference. For the original isotropic model, classical surrogates exist and NPE is primarily a computational convenience—it replaces per-observation ABC costs with a one-time training investment. For Model A (directional bias) and Model B (proliferation), surrogates become increasingly approximate, and NPE offers both computational and accuracy advantages by avoiding mean-field approximations. For Model C (combined bias and proliferation), the interplay of multiple mechanisms renders mean-field surrogates difficult to validate, and NPE provides a principled route to full posterior inference by bypassing the surrogate entirely. This progression underscores NPE’s role as a general-purpose inference engine that scales naturally with model complexity, performing well across all four models regardless of whether a reliable surrogate happens to exist.

NPE is one of several neural approaches to simulation-based inference [21, 43]. Neural Ratio Estimation (NRE) learns the likelihood-to-evidence ratio and requires MCMC sampling to obtain posterior samples [51], while Neural Likelihood Estimation (NLE) learns a surrogate likelihood function that must likewise be combined with MCMC [52]. We chose NPE because it directly produces the posterior density without any additional sampling step, yielding a self-contained inference procedure that requires no MCMC post-processing. Benchmark studies in low-to-moderate-dimensional problems show that NPE performs competitively with NRE and NLE in terms of posterior accuracy [53]. As a practical bonus, the learned posterior is amortized: once trained, each new observation yields posterior samples via a single forward pass, which is advantageous when analysing multiple experimental replicates. We note that NRE or NLE may be preferable for problems where the likelihood-to-evidence ratio is simpler to learn than the full posterior, or where sequential refinement around a single observation is prioritised over amortization across many observations.

### 4.2 Role of Spatial Information

Our results reveal a nuanced picture of when spatial information improves inference. For the original isotropic model, 1D column counts and 2D spatial data yield comparable posterior precision (Table 5), confirming that well-chosen summary statistics are effective when the model’s symmetry aligns with the summary statistic. This framing clarifies the role of each data representation: 1D column counts are hand-crafted summary statistics that exploit known model symmetries, while the 2D CNN performs automated feature extraction directly from raw spatial data. For isotropic models, the CNN learns features functionally equivalent to column counts; the two approaches converge because no additional spatial information exists to exploit.

The value of spatial inference is demonstrated concretely by the extended models. Model A (directional bias) creates anisotropic spreading patterns where agents preferentially migrate in one direction, producing spatial asymmetries that are collapsed when column counts sum across rows. The 2D CNN has access to these directional features, and while the *P* –*ρ* degeneracy persists in both representations (Section 3.2), the spatial approach provides a representation that naturally accommodates anisotropy without requiring the analyst to design bias-aware summary statistics. When comparing matched training budgets (50,000 simulations for both 1D and 2D), the benefits of spatial data are parameter-dependent. In Model A, the 2D posterior for *ρ* is approximately 1.7 times tighter than 1D ( ∼0.59*×* the CI width), reflecting the spatial asymmetries created by directional bias, while *P* is comparably constrained by both representations. In Model B, the 1D posteriors are at least as tight as the 2D posteriors for all three parameters, indicating that column counts already capture the information relevant to this proliferation model. In Model C, the picture is mixed: the 2D posteriors are tighter for *P* ( ∼0.86 times) and *ρ* ( ∼0.70 times), while *U* and *R* are comparably or better constrained by 1D column counts, suggesting that spatial information provides additional constraining power for some parameters but not uniformly across all.

More broadly, the cost-benefit calculation shifts decisively toward spatial approaches for models where crucial information is encoded in patterns that resist simple summarization: agent-agent interactions, chemotaxis, cell-cell adhesion, heterogeneous environments, and multi-species dynamics all generate spatial structure that 1D projections discard. When the degenerate structure is known *a priori*, reparameterizing to identifiable continuum parameters (Section 3.2) provides an additional route to improved calibration beyond increasing the training budget or changing the data representation.

Interpretability remains an open challenge for CNN-based inference. Techniques such as activation mapping [54] or saliency maps [55] could visualize the spatial features our CNN learns to associate with different parameter values. The network likely develops sensitivity to biologically relevant patterns like the sharpness of the spreading front, local cell density gradients, or the degree of cell clustering. By identifying these learned features, a researcher could validate that the network’s reasoning aligns with established biological theory, or discover novel spatial statistics that are informative but not immediately obvious to a human observer. While a detailed feature analysis is beyond the scope of this work, we highlight its importance for future applications. The ability to not only perform inference but also to extract the learned representations that enable it is a distinct advantage of this approach.

The ability to perform inference with stochastic models on spatiotemporal data has direct implications for experimental design. Traditional protocols often reduce spatiotemporal data from time-lapse microscopy to simplified measurements, largely because of past computational limits. Our framework removes this barrier, enabling direct analysis that preserves the full spatial information. This should encourage the strategic collection and retention of rich datasets, which is crucial for improving parameter identifiability in complex systems where spatial structure encodes mechanistic information.

### 4.3 Computational Scalability and Future Directions

The primary cost for any NPE approach is the upfront simulation budget, which can become substantial for models with high-dimensional parameter spaces (*>* 15–20 parameters). On top of this, the CNN adds specific challenges: (i) memory requirements scale quadratically with spatial resolution, potentially limiting application to very high-resolution imaging data without careful optimization; (ii) architecture selection for CNN embedding networks requires problem-specific tuning of depth, width, and kernel sizes, adding complexity to the analysis pipeline; (iii) the combined optimization of feature extraction and density estimation can be more challenging than simpler approaches, requiring careful hyper-parameter selection. (We note that for the lattice sizes considered here, CNN training adds negligible overhead; this concern applies primarily to higher-resolution spatial data.)

These costs are decreasing: GPU-accelerated simulators can reduce the simulation bottleneck by orders of magnitude, improved flow architectures reduce the training budget needed for accurate posteriors, and transfer learning may allow pre-trained spatial feature extractors to be adapted to new biological problems without retraining from scratch. Natural extensions of this framework include incorporating temporal dynamics from time-lapse microscopy (multiple snapshots rather than a single time point), three-dimensional spatial data from confocal imaging, and multi-species models where inter-population interactions generate spatial structure that resists simple summarization.

### 4.4 Conclusions

We have demonstrated that NPE provides a principled path beyond the long-standing tension between computational tractability and biological realism in parameter inference for stochastic agent-based models. Across a hierarchy of random walk models—from isotropic diffusion through to a four-parameter model combining directional bias and proliferation—NPE performs well regardless of whether a reliable surrogate happens to exist. Integrating CNNs as learned feature extractors enables inference directly from high-dimensional spatial data, removing reliance on hand-crafted summary statistics. We provide an open-source implementation of our complete pipeline^4^ to facilitate adoption and extension to more complex, spatially-structured biological models.

## A Posterior Predictive Checks

To complement the simulation-based calibration diagnostics presented in Section 3.5, we evaluate NPE’s predictive capabilities through posterior predictive sampling for the original model. For each posterior sample *θ*^(*j*)^, we generate predicted data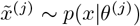using the stochastic simulator. The collection of predictions forms empirical prediction intervals that naturally account for both parameter uncertainty and model stochasticity.

### A.1 1D Column Count Predictions

As shown in Figure 10, the resulting NPE-based predictions are well-calibrated, with 94.8% of true observations falling within the 95% prediction intervals. A key advantage of this simulator-native approach is its physical consistency; the prediction intervals respect the non-negative, discrete nature of the count data without requiring an explicit, and potentially erroneous, noise model. The framework also naturally captures heteroscedastic uncertainty, with the prediction intervals appropriately widening in regions of high agent density where stochastic variability is greatest.

**Figure 10.**
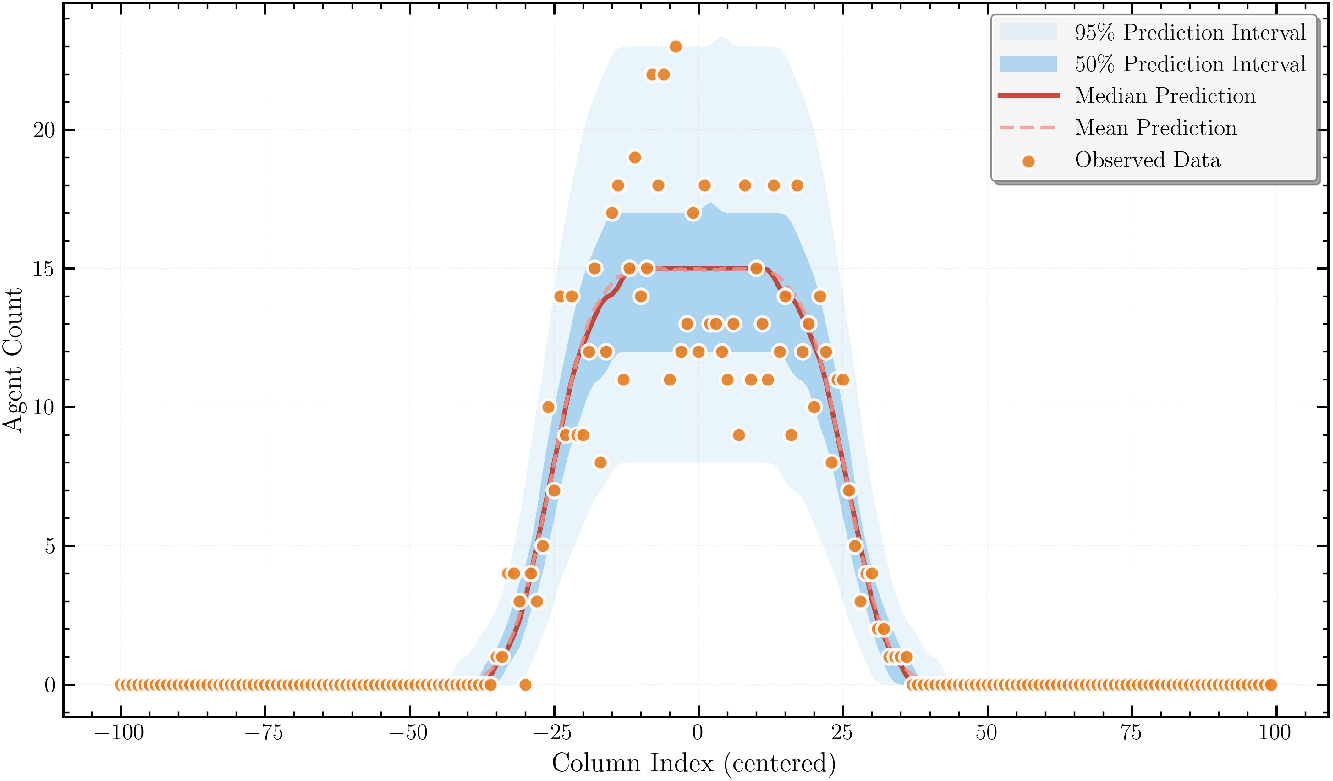
Posterior predictive checks from NPE using 1D column count data. The observed data (orange circles) is overlaid on predictions generated by simulating the model with parameters drawn from the inferred posterior distribution. The median (solid red line) and mean (dashed red line) of the predictions are shown, along with the 50% (dark blue) and 95% (light blue) prediction intervals. For visualization purposes, a Gaussian kernel has been applied to smooth the predictive distributions. An appropriate proportion of the observed data points lie within the 95% prediction interval, indicating that the calibrated model is consistent with the data.

### A.2 2D Spatial Predictions

As shown in Figure 11, the 2D-trained model generates predictions consistent with the observed data. The calibrated model accurately captures the spatial characteristics of the agent-based simulation, producing prediction intervals that are well-calibrated and physically consistent with the discrete nature of the data. Importantly, the framework captures the input-dependent (heteroscedastic) uncertainty, something that standard MLE under an independent and identically distributed noise assumption (used in much of the mathematical biology literature) cannot represent, often leading to miscalibrated, homoscedastic intervals even when the underlying variability is spatially varying. As in the 1D case, the method remains robust when applied to high-dimensional (2D) spatial inputs.

**Figure 11.**
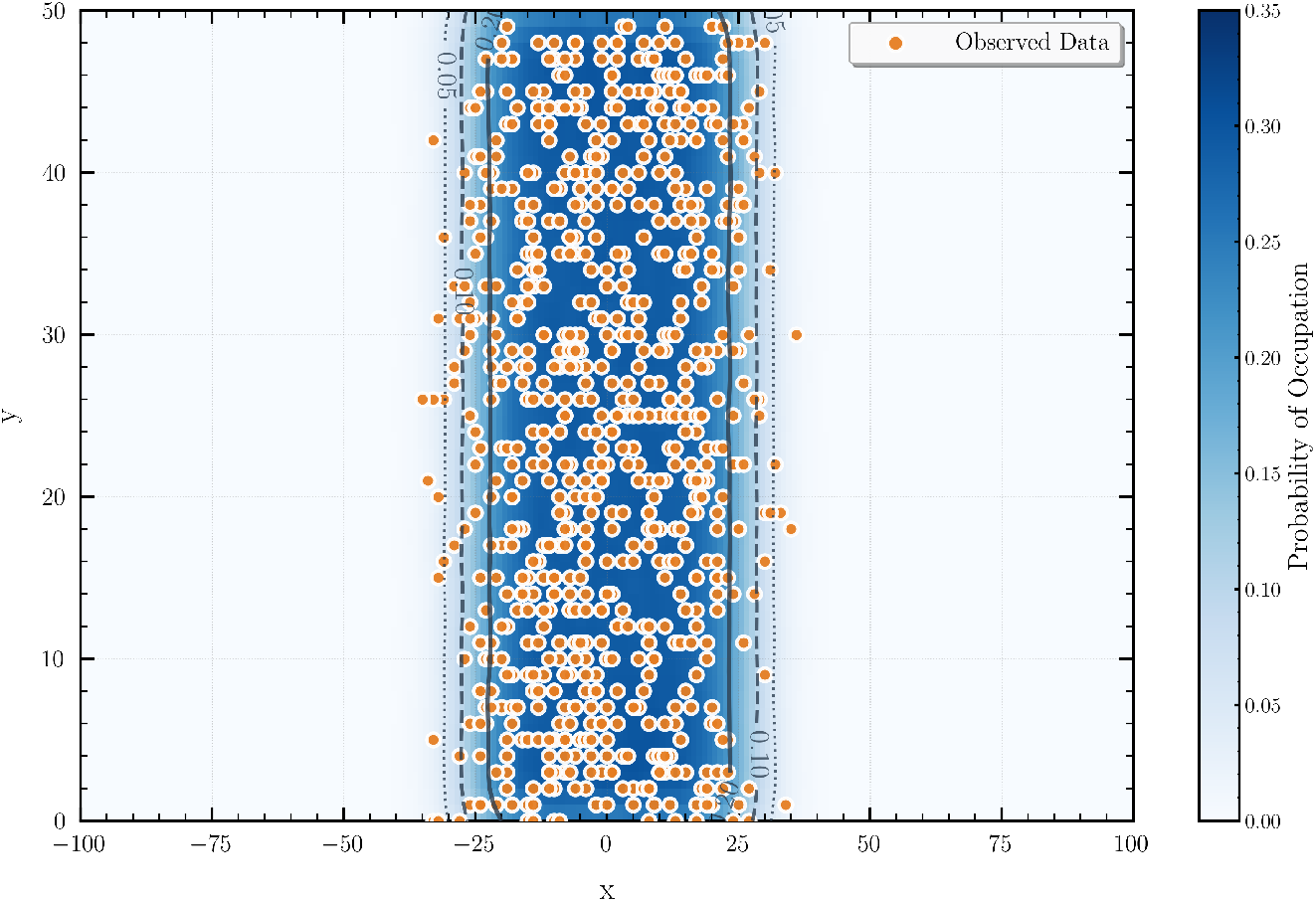
Posterior predictive check for the 2D spatial data model, showing the full observed data points (orange) overlaid on the predicted occupancy probability from the NPE model (blue heat-map). The grey contours delineate iso-probability curves at probability levels of 0.05, 0.10, and 0.20 (dotted, dashed, solid, respectively).

In the 2D (CNN) results we also note a modest separation between the posterior predictive mean and median, consistent with a mild right-skew in the posterior for *D*. We attribute the observed skew primarily to structural features of the inference problem—namely a slight *U* –*D* trade-off, support constraints (*D*≥ 0 with *D* = *P/*4), and the non-linear parameter-to-observable map—together with the fact that the CNN accesses richer spatial cues (e.g., front roughness and local clustering) that are averaged out in 1D summaries and therefore make such asymmetries more apparent. We cannot entirely rule out minor artefacts arising from the higher-dimensional learning task.

## B Reproducibility Across Random Seeds

Neural network training involves stochastic elements—random weight initialisation and mini-batch ordering—that could in principle cause posterior estimates to vary between independent runs. To quantify this variability, we retrained the 1D NPE pipeline five times for each model using different random seeds (42, 123, 456, 789, 1024) while holding the test observation fixed. Table 7 summarises the results. Posterior mean estimates are stable across seeds: the standard deviation of the posterior mean is at most 0.032 (*P* in Model A), which is small relative to both the posterior width and the parameter range. Credible interval widths are similarly consistent, with relative fluctuations typically below 10%. The true parameter value falls within the 95% credible interval for all five seeds and every parameter (5/5 coverage), though we note the small sample size limits the statistical power of this check. These results confirm that the trained posteriors are robust to the stochastic elements of the training procedure and that a single training run is sufficient for reliable inference. We observe similar stability for the 2D NPE posteriors across seeds.

**Table 7:**
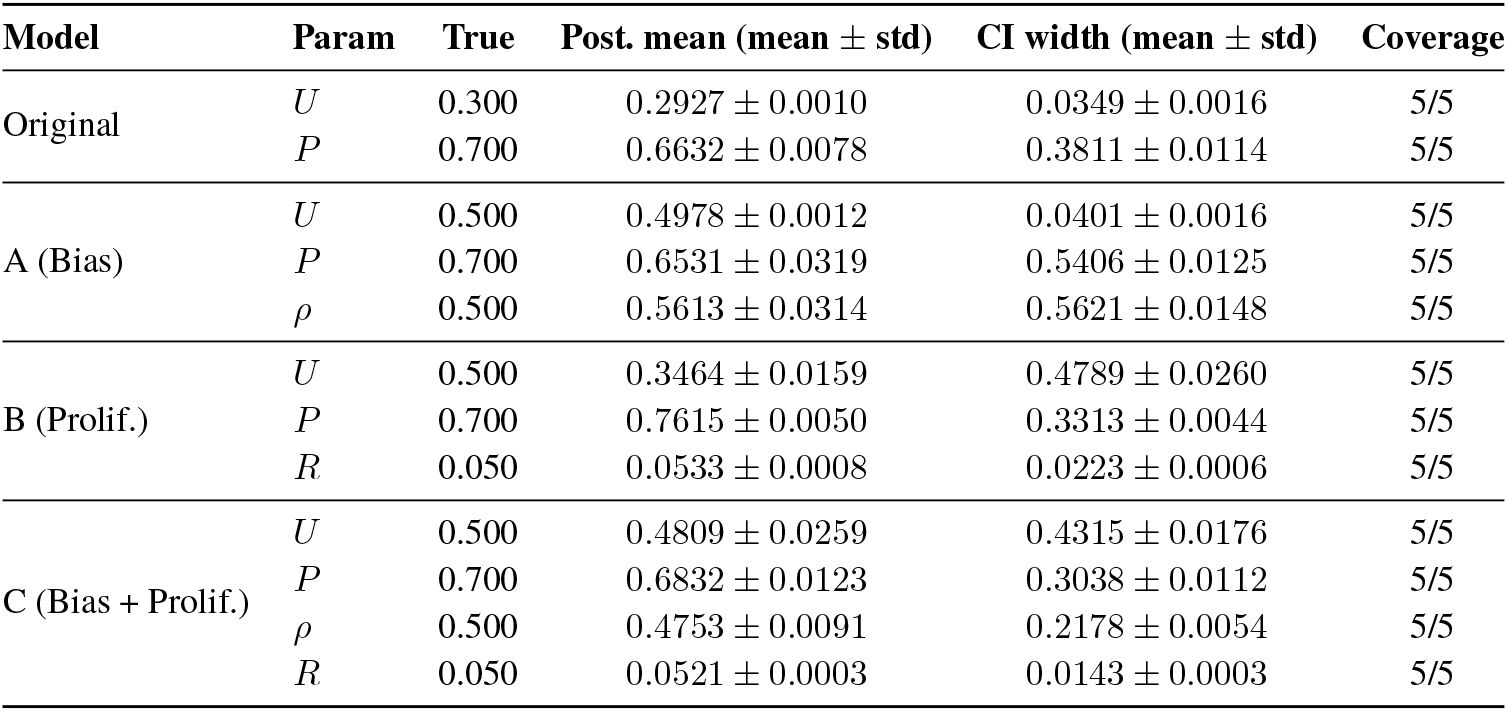
Reproducibility across random seeds. For each model and parameter, we report the mean ± standard deviation of the posterior mean and the 95% credible interval width across 5 independent training runs. Coverage indicates the fraction of seeds for which the true value falls within the 95% credible interval. All 1D NPE was performed with 50,000 training simulations.

## Footnotes

2 https://github.com/tomkimpson/NPE_for_RW_inference

3 Although parameter identifiability is not a significant concern for the simple model we consider, additional spatial features are of interest for disentangling parameters in more complex models that incorporate processes like cell-cell adhesion or proliferation.

4 github.com/tomkimpson/NPE_for_RW_inference

